# Estimation of the force of infection and infectious period of skin sores in remote Australian communities using interval-censored data

**DOI:** 10.1101/674663

**Authors:** Michael J. Lydeamore, Patricia T. Campbell, David J. Price, Yue Wu, Adrian J. Marcato, Will Cuningham, Jonathan R. Carapetis, Ross M. Andrews, Malcolm I. McDonald, Jodie McVernon, Steven Y. C. Tong, James M. McCaw

## Abstract

Prevalence of impetigo (skin sores) remains high in remote Australian Aboriginal communities, Fiji, and other areas of socio-economic disadvantage. Skin sore infections, driven primarily in these settings by Group A *Streptococcus* (GAS) contribute substantially to the disease burden in these areas. Despite this, estimates for the force of infection, infectious period and basic reproductive ratio — all necessary for the construction of dynamic transmission models — have not been obtained. By utilising three datasets each containing longitudinal infection information on individuals, we estimate each of these epidemiologically important parameters. With an eye to future study design, we also quantify the optimal sampling intervals for obtaining information about these parameters. We verify the estimation method through a simulation estimation study, and test each dataset to ensure suitability to the estimation method. We find that the force of infection differs by population prevalence, and the infectious period is estimated to be between 12 and 20 days. We also find that optimal sampling interval depends on setting, with an optimal sampling interval between 9 and 11 days in a high prevalence setting, and 21 and 27 days for a lower prevalence setting. These estimates unlock future model-based investigations on the transmission dynamics of GAS and skin sores.

## 1 Introduction

Infections with impetigo (commonly known as skin sores) remain highly prevalent in remote Australian Aboriginal communities, as well as Fiji and areas of socio-economic disadvantage [3, 36]. Skin sore infections in these settings are primarily caused by *Staphylococcus aureus*, and Group A *Streptococcus* (GAS). GAS is associated with post-infectious sequelae such as acute rheumatic fever and rheumatic heart disease, of which Australia has one of the highest recorded prevalences globally [3]. Despite a relatively high level of understanding about the specifics of the GAS bacterium [39, 30, 34, 9, 16], comparatively little is known about the natural history of skin sore infection. Furthermore, what is known is often based on historical studies from a generation prior and from a different, non-endemic, geographical region [12, 10, 24, 11]. We aim to utilise a dynamic transmission model for skin sores to estimate two key quantities: the *force of infection*, and the *duration of infectiousness*. In the absence of information relating to immunity post-infection, we assume skin sore transmission follows the dynamics of the Susceptible-Infectious-Susceptible (SIS) model. Calculation of these two key quantities will contribute to the development and parameterisation of models which will in turn inform the design of intervention strategies aimed at reducing prevalence.

We analyse three separate datasets, all from remote Australian communities, documenting the infection dynamics of skin sores in individuals. The first dataset consists of public health network presentation data for 404 children under five years of age [19, 8, 26], collected as part of the East Arnhem Healthy Skin Project; the second contains longitudinal data for 844 individuals from three rural Australian communities, collected during household visits [25], and the third is comprised of survey visits for 163 individuals who participated in a mass treatment program [5], of which the primary endpoint was control of scabies infection. To analyse these data, we linearise the SIS model about the endemic equilibrium, and derive an expression for the likelihood of the two model parameters. By utilising Markov chain Monte Carlo (MCMC) methods, we obtain estimates of each of the force of infection, the duration of infectiousness, the basic reproductive ratio, *R*_0_, and the prevalence of infection. Finally, by utilising optimal experimental design, the optimal sampling strategy to inform estimation of these parameters for use in future studies is obtained.

## 2 The Susceptible—Infected—Susceptible model

We consider a stochastic representation of the Susceptible-Infectious-Susceptible (SIS) model [22]. In this model, individuals are either susceptible (*S*) or infectious (*I*). The transition rate from susceptible to infectious, known as the *force of infection*, and denoted λ = *βI*, where *β* is the transmissibility parameter, is non-linear. This non-linearity is one of the key features of dynamic infectious disease models. However, this means that to model a population of individuals, the state of each individual is required (to know the prevalence, *I*). For the SIS model, the size of the required state space is *N* [21]. When constructing the Markov chain representation of the SIS model then, the generator matrix, *Q*, is *N* × *N*, meaning that for large numbers of individuals, computing the matrix exponential exp(*Qt*), is computationally intractable. The result of this, then, is that performing inference with infectious disease models is challenging [29, 2, 27, 23, 40, 28].

When the dynamics of the SIS model are at (or close to) equilibrium, then the force of infection, λ, is approximately constant. As such, we approximate the SIS model by a two-state process with a *constant* force of infection. By making this approximation, and assuming individuals are otherwise identical, it follows that a Markov chain consisting of only two states is required, independent of the underlying population size. This approximation has a straightforward likelihood calculation, which allows estimation in a Bayesian framework and also the calculation of the optimal sampling interval for future study designs through the use of optimal experimental design.

### 2.1 Linearisation of the SIS model

The standard SIS model can be described using two transitions, infection and recovery, and two parameters, the transmissibility parameter, *β*, and the rate of recovery, *γ* (Table 1). Ignoring demographic processes, the total number of individuals in the population is fixed.

**Table 1:** Transitions of the SIS model. The force of infection is given by λ, the transmisibility parameter β and the rate of recovery by *γ*.

| State | $\Delta$ | Rate |
| --- | --- | --- |
| $(S, I)$ | $(-1, +1)$ | $\lambda := \beta I$ |
| $(S, I)$ | $(+1, -1)$ | $\gamma$ |

One of the most important quantities in infectious disease modelling is the *basic reproductive ratio, R*_0_, defined as the mean number of secondary infection events caused by a single infectious host, in an otherwise susceptible population. The basic reproductive ratio functions as a threshold parameter, where if *R*_0_ ≤ 1, an outbreak of disease will not occur, while if *R*_0_ > 1, then there is a non-zero probability of a disease outbreak occurring. For the SIS model, the basic reproductive ratio is

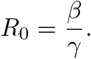

The quasi-equilibrium solution of the SIS model is well known [21], and the endemic prevalence of disease is

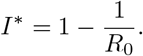

Let *t* be time of interest of the process. The force of infection, λ(*t*) is defined as,

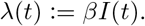

At equilibrium, the prevalence is approximately constant, and so the force of infection can be approximated by

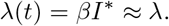

By performing this linearisation, it is assumed that the dynamics of disease are and remain at equilibrium. It follows that we may consider a single individual. The generator matrix for the Markov chain for the life-course of that individual is

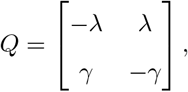

and the matrix exponential of *Q* is

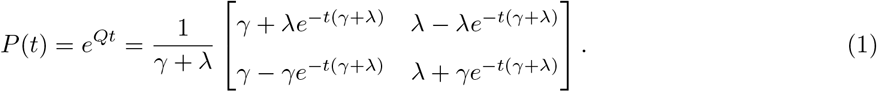

The matrix in Equation (1), combined with an initial state and time *t*, gives the probability distribution for the Markov chain. It is possible to calculate expressions for the equilibrium prevalence, *I*^*^, and the basic reproductive ratio, *R*_0_, in terms of λ and *γ*. Solving for the equilibrium distribution of the linearised SIS model

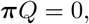

gives the equilibrium prevalence

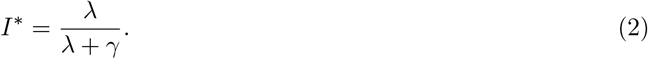

From the standard SIS model, it is also known that the basic reproductive ratio, *R*_0_ = *β*/*γ*, and λ = *βI*^*^. It follows that the basic reproductive ratio, *R*_0_, is given by

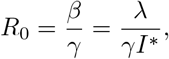

and substituting Equation (2) gives

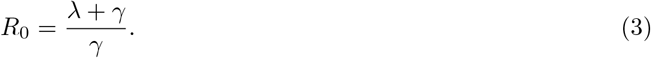

Given these simple closed form expressions for the key quantities of interest, it is possible to perform estimation in a Bayesian setting, using interval-censored data.

## 3 Data

Three separate datasets collected in Australian Aboriginal communities are considered in this study: data from public health network presentation (PHN) data on 404 children from birth to five years of age, collected as part of the East Arnhem Healthy Skin Project; data for 844 individuals from three communities, collected during household visits (referred to as the HH dataset); and data from 163 individuals who were observed for over 25 months as part of a mass treatment program in a single rural community (RC). Ethics approval for reuse of existing data was obtained from The Human Research Ethics Committee of the Northern Territory Department of Health and Community Services and Menzies School of Health Research (Ethics approval number 2015-2516). Permission was also obtained from the custodians of each dataset. This project has been conducted in association with an Indigenous Reference Group, as well as an ongoing stakeholder group which contains Aboriginal Australian community members. Each dataset consists of longitudinal observations of each individual, where their infection status is recorded at each observation. The times between presentations are heavily right skewed in each dataset, with a median time to next presentation of 9 days for the PHN data, 61 days for the HH data and 119 days for the RC data. The number of observations in total is also highly variable with 13,439 observations in the PHN data, 4,507 in the HH data and 626 in the RC data. Kernel density estimates of the distribution of time until the next presentation, with the observed data overlayed, are shown in Figure 1. The suitability of each of these datasets for inferring the force of infection, λ, rate of recovery, *γ*, and the basic reproductive ratio, *R*_0_ is investigated in Section 4.3. It is worth noting specifically that the PHN data contains information only on children from birth to five years of age, while the other two datasets contain information on individuals of all ages. Prevalence of skin sores is known to be age-dependent [35] and so by not modelling any age-structure, we are ignoring these differences.

**Figure 1:**
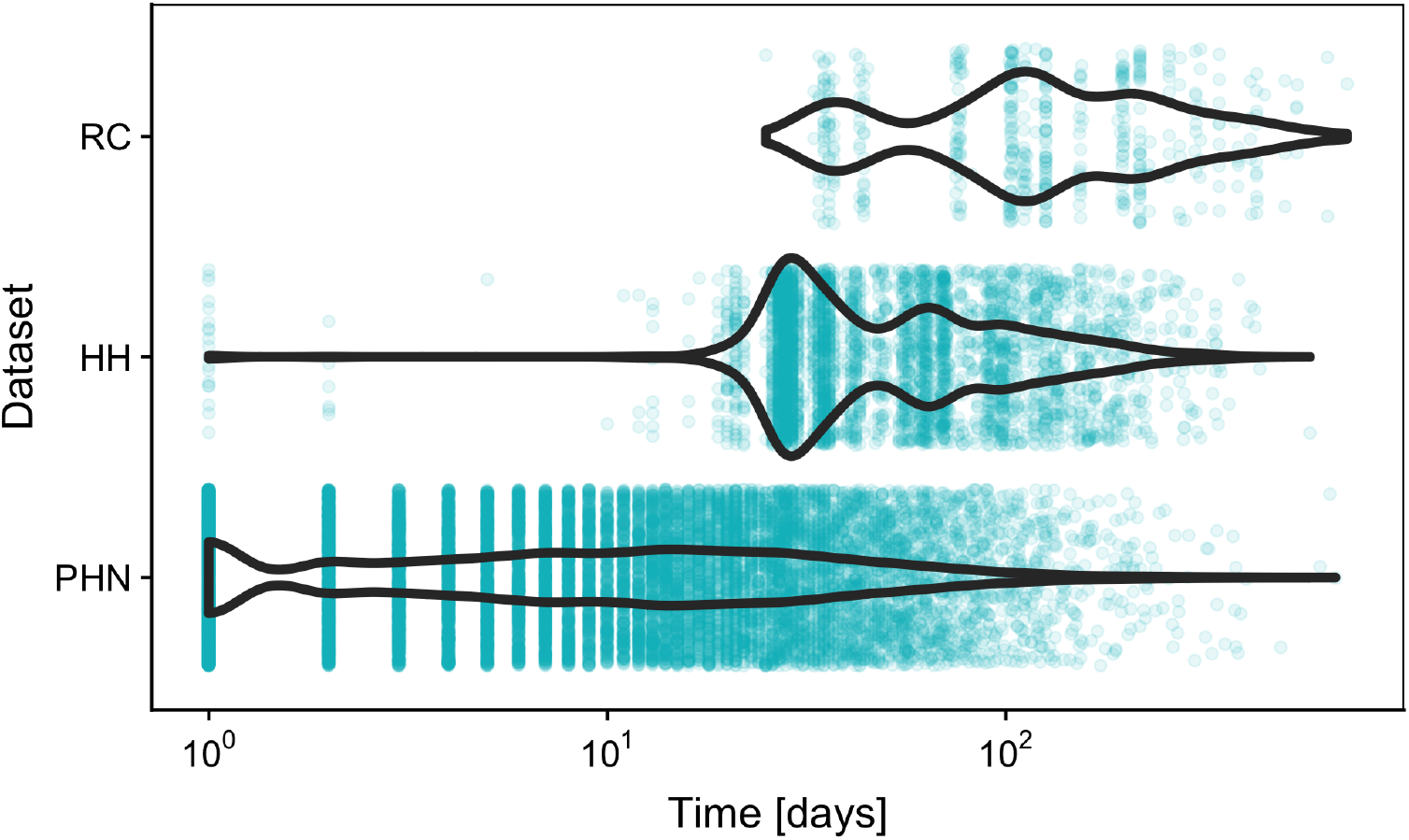
Distribution of time between presentations for each of the three datasets — PHN, HH and RC — with the time between observations overlayed.

### 3.1 Data Structure

Recall that the datasets which are considered consist of longitudinal observations for each individual, with an individual’s infection status being noted as either susceptible or infected at each point. The observation is not continuous in nature, with the individual’s infection status only being known at each sampling point. Data of this form are known as *interval-censored*, or *panel* data. Interval-censored data are common in epidemiology, and inference in a frequentist setting is well established [18]. Let the state of individual *i* at observation *j* be *X_i,j_*, and the time at which the *j*th observation is made be *t_i,j_*. The likelihood for a single individual, *i*, can be evaluated as

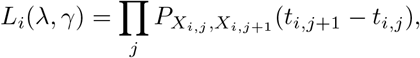

which is the relevant entry of the *P* matrix in Equation (1), evaluated at the time difference between observations, *t*_*i,j*+*i*_ – *t_i,j_*. It follows that the likelihood for the entire population is

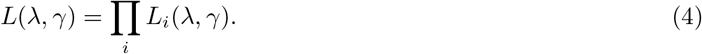

It is important to note that the likelihood in Equation (4) has assumed what is known as *ignorable* sampling times. That is, the sampling times are chosen independently of the outcome of the process. When sampling times are chosen in advance, as they were in the HH and RC datasets, then the sampling times have been proven to be ignorable [18]. For the PHN data, observations were made under what is termed a *doctor’s care* scheme, whereby the next observation time is chosen at the current observation time, and based on an individual’s disease state at that time. The sampling times are proven to be ignorable if the following two conditions are true [15]:

- The probability of individual i being in a given disease state *u_i,j_* at time *t_i,j_*, given all infection history until this point, *H*_*i,j*−1_, is independent of whether an examination is carried out at this time and past examination times, and
- The conditional distribution of the *j*th observation time, *t_i,j_* = *P*(*T_i,j_* = *t_i,j_*|*H*_*i,j*−1_), where *T_i,j_* is the random variable representing the time of the *j*th infection for individual *i*, is functionally independent of the transmission parameters.

The first of these conditions effectively means that the probability that an individual is either susceptible or infectious at time *t_j_*, given all past information, is independent of *t_j_*, and all past examinations. As treatment is prescribed by a doctor’s visit, it is possible that this condition is violated. However, the dataset does not contain information on the form of treatment administered meaning that it cannot be assumed that the administered treatment is for skin sores, and almost 60% of presentations to the clinic contain no information on skin sores (and so one could assume that the primary reason for the visit is not skin sores). Further, it is noted that the estimate of infectious period in any modern setting will be *augmented* by treatment. As such, it is assumed that the first condition is true. The second condition means that the next observation time is conditionally independent of the transmission process. This condition is assumed to be true here due to the high frequency of presentation in this dataset, even when an individual does not have skin sores. It is important to note that it has been proven *impossible* to test whether or not a sampling scheme is ignorable. The analysis proceeds on the basis that all sampling schemes in the given data are ignorable, but it is noted that this may not be the case.

Both the force of infection, λ, and the infectious period, *γ* are estimated in a Bayesian context using Markov chain Monte Carlo estimation (MCMC). The MCMC is performed using the No-U-Turn sampler implemented in Stan [37], using 10,000 iterations for 4 chains, for a total of 40,000 iterations. The code used to perform this estimation is available at http://github.com/MikeLydeamore/TMI/.

## 4 Analysis

We first start by verifying the methodology through the use of a simulation estimation study, whereby individuals are simulated from the linearised SIS model, and attempt to recover parameters through the estimation routine detailed in the previous section. Then, the estimation method is applied to the observational data.

### 4.1 Verification of methodology

There are multiple sources of stochastic variability in this setting, including the underlying population which is observed, the realisation from the observation distribution and the MCMC method itself. The first two of these potential causes for variation are investigated in detail here.

To investigate the variability in the underlying population, the estimation procedure is performed on 64 randomly generated populations from the linearised SIS model, and each of the 400 members of each simulated population are observed once daily for one year. This high frequency of observation means that the only source of meaningful variability is that which comes from the linearised SIS model. The top row of Figure 2 shows the marginal posterior estimates for the force of infection, λ, the rate of recovery, *γ*, and the basic reproductive ratio, *R*_0_, for populations simulated using λ = 1/60 and *γ* = 1/20. Each violin plot shows an individual (marginal) posterior distribution for the parameter of interest from a randomly selected population, while the boxplot shows the variability of the posterior mean for each parameter over all 64 realised populations. The within-simulation variability is relatively high, even in this case with daily observation. However, the method estimates each parameter well and in an unbiased manner.

**Figure 2:**
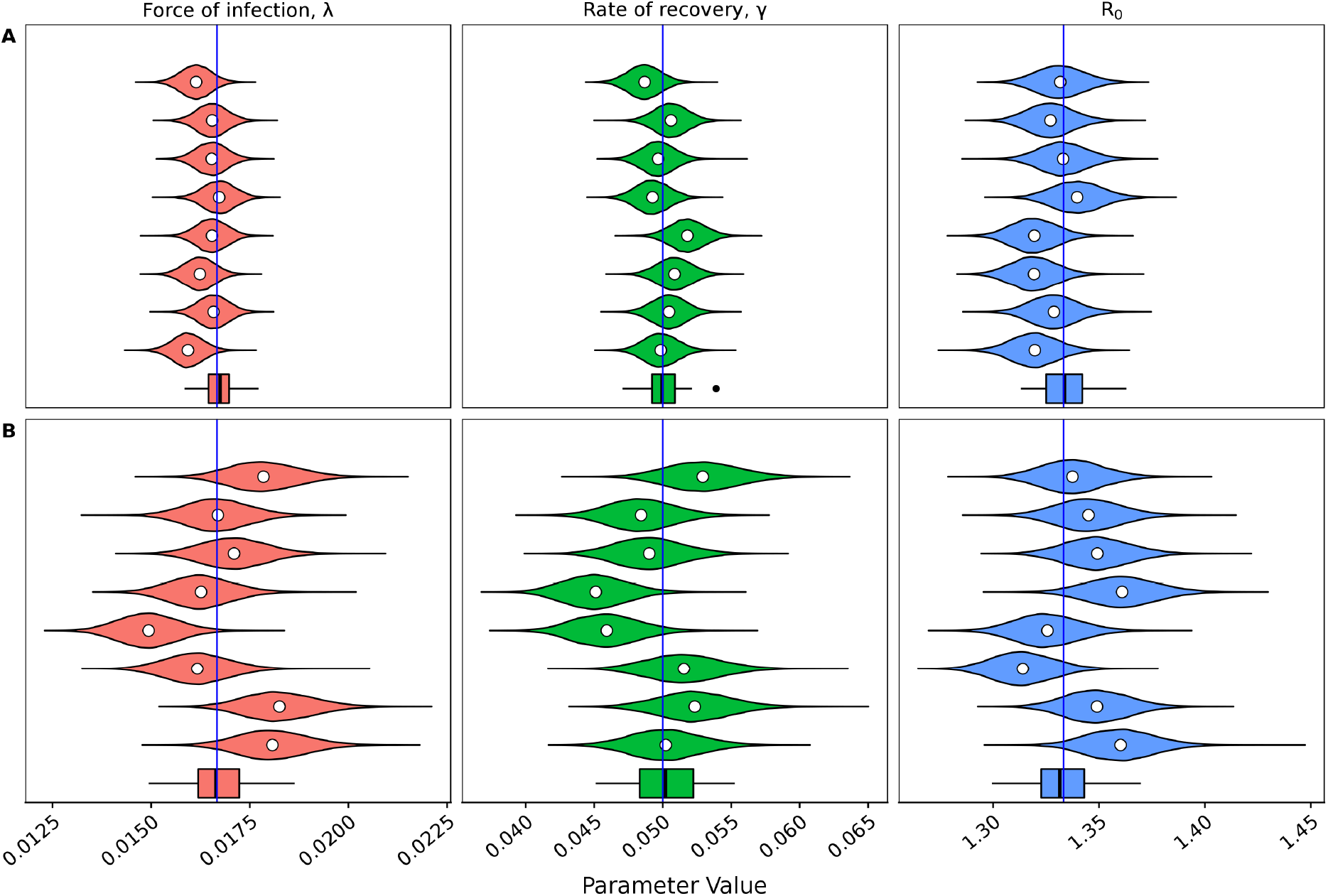
Marginal posterior distributions for the force of infection, λ, the rate of recovery *γ*, and the basic reproductive ratio, *R*_0_, from 8 randomly generated populations from the linearised SIS model under two different observation distributions. The mean of each distribution is given by the white circle. The boxplot at the bottom of each panel represents the means of 64 marginal posteriors. The true value which was used to generate each population is represented by the blue line (λ = 1/60, *γ* = 1/20). The two different observation distributions are (A): Observed daily over 1 year and (B) observed according to the empirical presentation distribution from the PHN data over 1 year. Both observation distributions yield good estimates of the simulated parameters. The observation distribution from the RC dataset was tested, but has not been visualised as the estimates were far from the true values (See Figure 3).

Next, potential variability in the observation distribution is considered. Again, a population of 400 individuals is simulated, and each simulated individual observed at times drawn from the observation distribution obtained from the PHN dataset (shown in Figure 1) over a time horizon of 1 year (Figure 2(B)). It is satisfying that although the sampling interval in the PHN dataset is notably longer than the daily case shown in Figure 2(A), the estimation method is still able to recover the simulated parameters. This suggests that oversampling the population gives little benefit to estimates of the parameters. Comparatively, it makes sense that if the sampling interval is too large, then no information will be gained. An example of this phenomenon is shown in Figure 3, where 20 samples are made of the population, separated by some sampling interval. The figure shows that a short sampling interval and a relatively short time horizon means that information about the parameters is difficult to recover. Similarly, a long sampling interval increases the variance in the parameter estimates. This suggests that there exists some *optimal* sampling interval. This concept will be returned to in Section 5.

**Figure 3:**
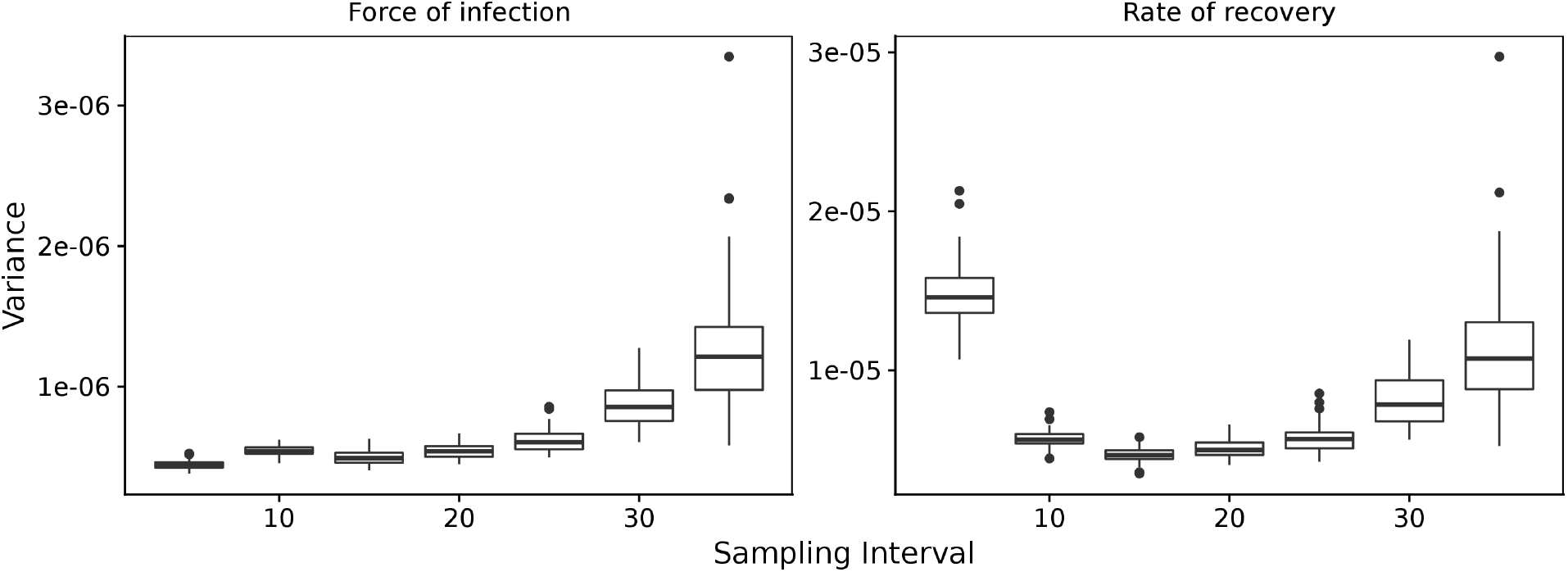
Variance in the estimates of the force of infection, λ, and the rate of recovery, *γ*, for a range of sampling intervals. Estimates were performed on 64 realisations of the simulated populations, each with parameters λ =1/60 and *γ* = 1/20. Each realisation contains 20 observations from the simulated population.

### 4.2 Verification of linearisation procedure

Having established that parameters can be re-estimated from the linearised model, we now look to verify whether the linearisation of the SIS model is valid. To do this, an individual-based implementation of the *full* (non-linearised) SIS model is used. The chosen parameters are *β* = 0.067 and *γ* = 1/20 (giving λ = 1/60 and an endemic prevalence of 25%), and 300 individuals. The Markov chain is seeded with 125 infected individuals, which is close to the equilibrium of this system. The population is simulated from the full SIS model, and the estimation is performed using the *linearised* model. Figure 4 shows results from 64 realised populations, under the observation distribution from the PHN dataset. The recovery rate, *γ*, is estimated accurately and with relatively small variance. The force of infection, λ, is somewhat underestimated on average with a relative error in the mean of 15%, although the variability is large. This underestimate carries over to the estimate of the basic reproductive ratio, *R*_0_.

**Figure 4:**
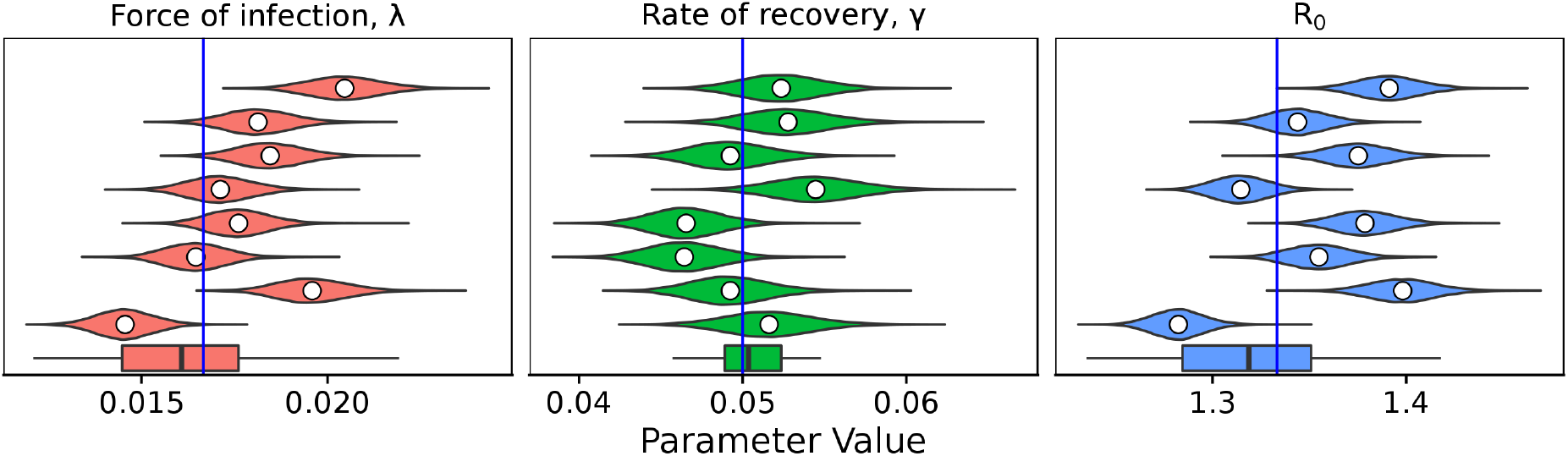
Marginal posterior distributions for the force of infection, λ, the rate of recovery *γ*, and the basic reproductive ratio, *R*_0_, from 8 randomly generated populations from the full (nonlinearised) SIS model under the empirical observation distribution from the PHN data, over 1 year. The mean of each distribution is given by the white circle. The boxplot at the bottom of each panel represents the means of 64 marginal posteriors. The true value which was used to generate each population is represented by the blue line (λ = 1/60, *γ* = 1/20). The simulated parameters are recovered successfully.

The results are visually similar to those shown in Figure 2 under the same observation distribution. Thus, it is concluded linearisation of the SIS model is valid when the dynamics are near equilibrium.

### 4.3 Verification of presentation distributions

Before estimating the force of infection, λ, and the rate of recovery, *γ*, for the three datasets discussed, the frequency of presentations must be checked to determine if they are sufficient for use with the method. Figure 5 shows a simulation estimation study using the presentation distributions from the PHN and HH datasets. Both datasets give good estimates. When considering the RC dataset, recall the presentation distributions shown in Figure 1. The RC dataset has a much wider sampling interval compared to the PHN and HH datasets. It is suspected that this presentation distribution may not hold sufficient information to recover the parameters of interest. However, as the prevalence is observed at each survey visit, estimating the basic reproductive ratio, *R*_0_, may still be possible. Figure 6 shows the prior distributions, with samples from the posterior distribution overlayed, under the observation distribution from the PHN dataset (panel A) and the RC dataset (panel B). Under the observation distribution from the PHN data, the posterior distribution samples are tightly clustered, with variance much smaller than in the prior distributions. Indeed, estimates are so localised relative to the prior that the samples appear to be overlayed in the figure. Comparatively, when the observation distribution is that seen in the RC dataset, the posterior samples are strongly correlated with a wide variance, indicating that this dataset does not have sufficient sampling frequency to separately estimate both the force of infection, λ, and the rate of recovery, *γ*. However, the posterior distribution samples align with the simulated prevalence (and thus the basic reproductive ratio, *R*_0_). The RC dataset can still be used to estimate these quantities.

**Figure 5:**
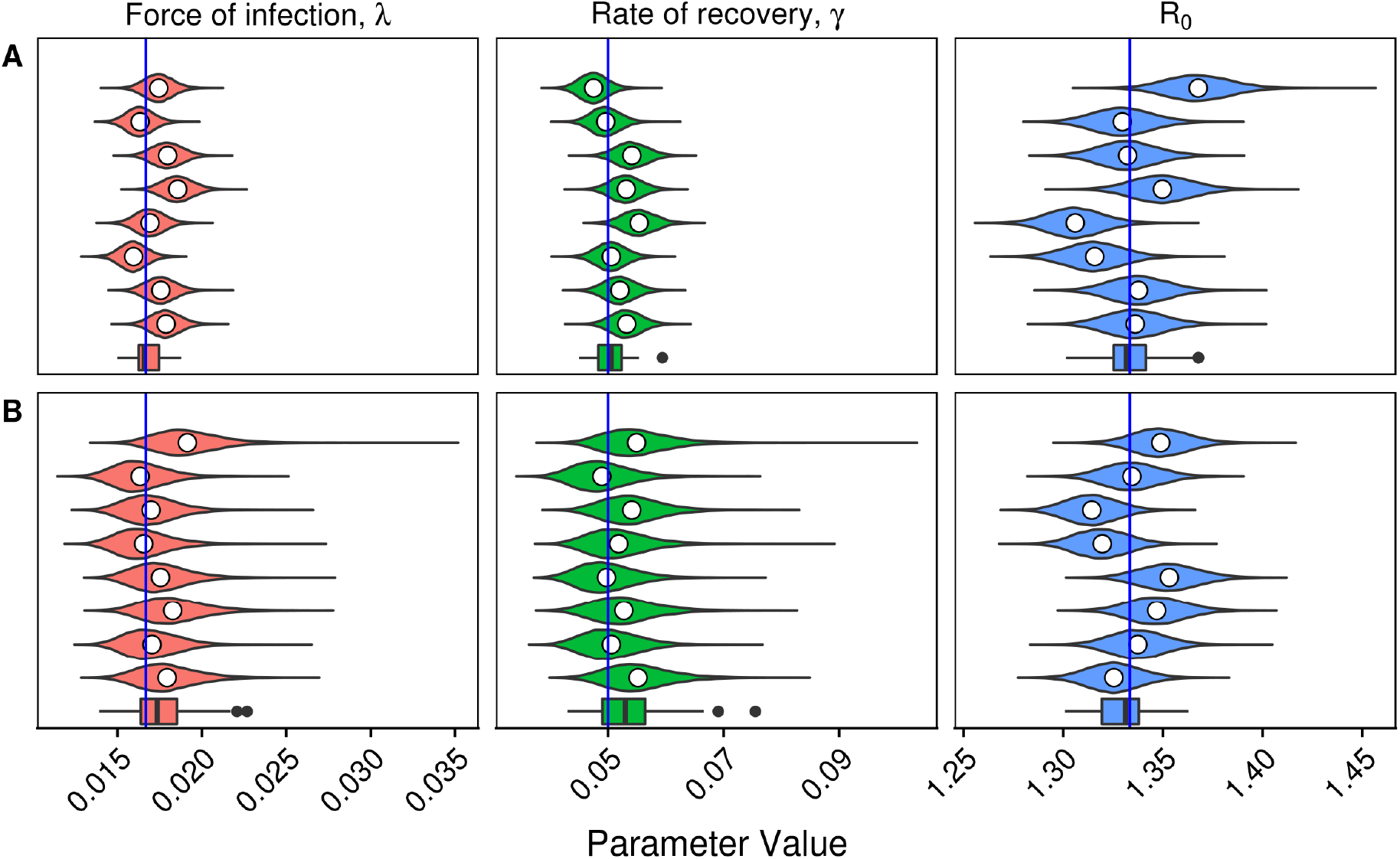
Marginal posteriors for the force of infection, λ, the rate of recovery *γ*, and the basic reproductive ratio, *R*_0_, from 8 randomly generated populations from the linearised SIS model under two different observation distributions. The mean of each distribution is given by the white circle. The boxplot at the bottom of each panel represents the means of 64 marginal posteriors. The true value which was used to generate each population is represented by the blue line (λ = 1/60, *γ* = 1/20). The two different observation distributions are taken from the (A): PHN dataset (1 year) and (B) HH dataset (1 year).

**Figure 6:**
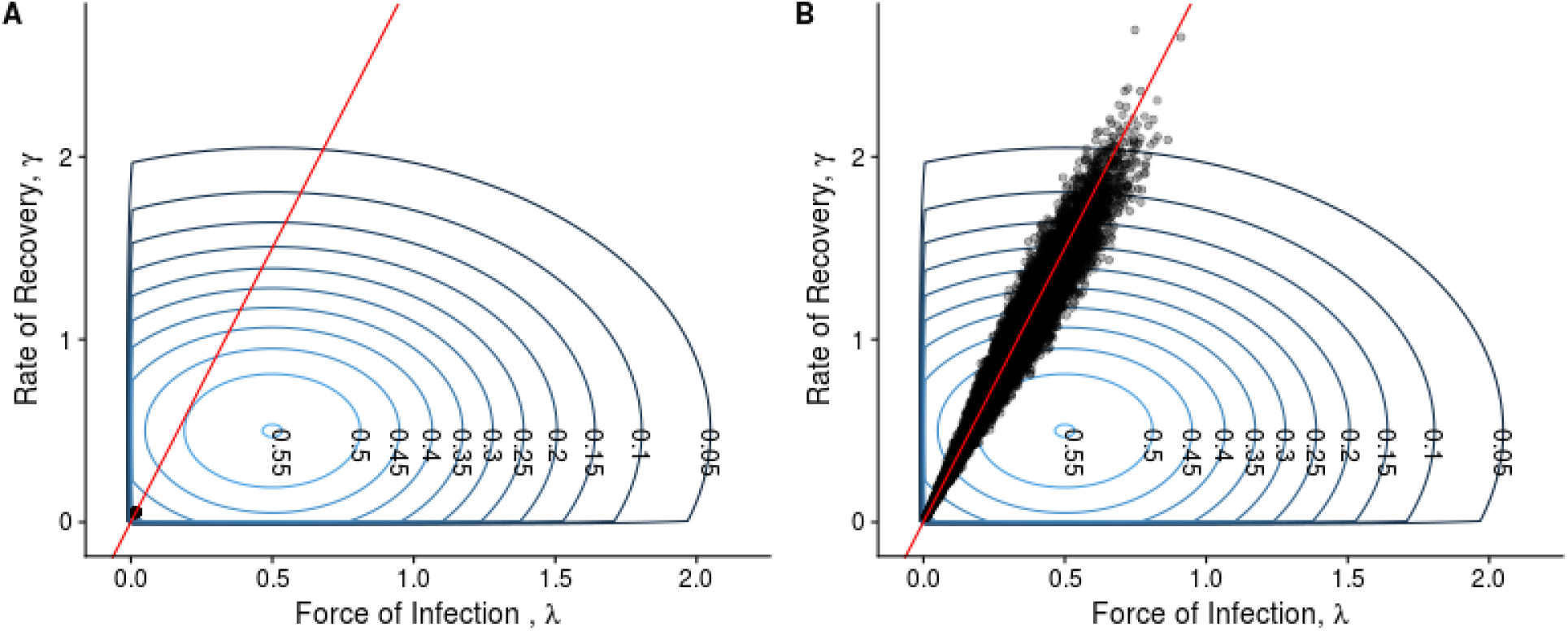
Prior distribution (concentric rings) with 20,000 samples from the posterior distribution (black points) overlayed from a randomly generated population under the observation distribution from (A) the PHN dataset, and (B) the RC dataset. The red line is the set of parameter values which give the true prevalence in the simulated population. In panel (A), the samples are tightly clustered with variance far smaller than the prior distribution. In panel (B), the samples are highly correlated, and with high variance, indicating the two parameters of interest cannot be uniquely determined, but their ratio (and so *R*_0_) can.

Having verified the suitability of each of the datasets to this estimation method, the next step is to estimate each of the force of infection, λ, the rate of recovery, *γ*, the prevalence of disease and the basic reproductive ratio, *R*_0_.

### 4.4 Estimation from Data

For the PHN and HH datasets, relatively similar estimates for the infectious period, 1/*γ* (12 days for the PHN dataset, and 20 days for the HH dataset) are obtained. However, notably different estimates for the force of infection, λ, were obtained. In the PHN dataset, the mean force of infection is estimated at 1/20.21, while in the HH dataset, the estimate is 1/202.07 — an order of magnitude different. This difference follows through to estimates of the basic reproductive ratio, *R*_0_ (1.60 vs 1.10), and the prevalence, estimated to be 37.5% in the PHN dataset and only 9% in the HH dataset. For the RC dataset, *R*_0_ is estimated to be 1.42, and the prevalence to be 26.9%. Point estimates of prevalence in all three study locations have been reported previously (Table 2)[1, 25, 5], at 35.6% in the region in which the PHN dataset was collected, 13.1% in the region where the HH dataset was collected and 35% in the region where the RC dataset was collected. These prevalence estimates align closely with the estimates obtained using our method.

**Table 2:** Parameter estimates for the force of infection, λ, and the infectious period, 1/*γ* from the three different datasets. Note this method estimates the rate of recovery, *γ*, but the infectious period is reported here for clarity.

| Dataset | Parameter [units] | Mean | 95% CI |
| --- | --- | --- | --- |
| PHN | Force of infection ($\lambda$) [1/days] | 0.049 | (0.042, 0.059) |
| | Infectious period ($\gamma^{-1}$) [days] | 12.19 | (10.23, 14.55) |
| | $R_0$ | 1.60 | (1.56, 1.65) |
|  | Prevalence | 37.5% | (31.0, 39.4) |
| Literature [1] | Prevalence | 35.6% | (32.9, 38.3) |
| HH | Force of infection ($\lambda$) [1/days] | 0.0049 | (0.0040, 0.0062) |
| | Infectious period ($\gamma^{-1}$) [days] | 19.97 | (16.19, 24.56) |
| | $R_0$ | 1.10 | (1.09, 1.11) |
|  | Prevalence | 9.1% | (8.3, 10.0) |
| Literature [25] | Prevalence | 13.1% | Not provided |
| RC | $R_0$ | 1.42 | (1.34, 1.51) |
| | Force of infection ($\lambda$) | — | Not identifiable |
| | Infectious period ($\gamma^{-1}$) | — | Not identifiable |
|  | Prevalence | 29.6% | (25.4, 33.8) |
| Literature [5] | Prevalence | 35% | Not provided |

## 5 Prospective Sampling Strategies

Thus far, the focus has been only on previously collected datasets from which to estimate parameters. If the sole aim of a study was to collect data to *best* estimate these parameters, then the natural question to ask is when should individuals be sampled? Aided by the simple structure of the linearised SIS model, this question may be answered through optimal experimental design [13]. We take the approach of robust optimal experimental design, under the ED-optimality criterion [31, 33]. Let ***δ*** = (*δ*_1_,…, *δ*_*n*−1_) define an n-sampling design with spacing *δ_i_, i* = 1,…, *n* – 1 between subsequent observations. Then, the optimal sampling design, ***δ***^*^, is given by

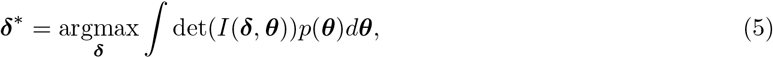

where ***θ*** = {λ, *γ*}, *I*(***δ, θ***) is the Fisher Information matrix, det is the determinant operator, and *p*(***θ***) is the prior distribution. Note that the optimal sampling interval, ***δ***^*^, is dependent on the prior distribution, *p*(***θ***). Two designs are considered for each dataset. The first is termed the *variable* sampling interval, where the ith sampling interval, *δ_i_*, is unrestricted, and *n* = 11 design spacings are chosen. Although this design strategy is optimal over a 12 visit design, adhering to the varying intervals may be difficult from an implementation perspective. A more practical strategy, and the second considered here, is termed the *fixed* sampling interval, where *δ_i_* = *δ*, ∀*i*. This is equivalent to considering *n* = 1 design spacing, as the population dynamics are assumed to be at equilibrium throughout the study.

The integral in Equation (5) is approximated using a Monte Carlo estimate with 5,000 samples. Each individual is observed 12 times. We use the induced natural selection heuristic for finding optimal strategies [32]. For detail on the algorithm inputs and evidence of convergence, see Appendix C.

### 5.1 Recommended sampling strategies

We calculate the optimal strategy using the posterior distributions obtained from the PHN dataset, HH dataset and the union of the these two posterior distributions as the prior distribution in Equation (5). The results for both the variable interval strategy and the fixed interval strategy are shown in Table 3.

**Table 3:**
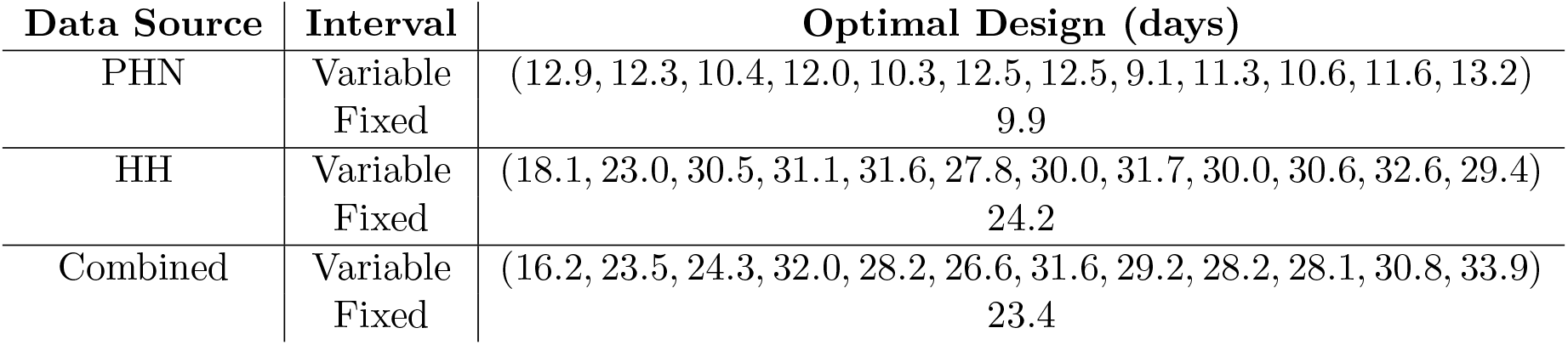
Optimal sampling strategies (in days) using the posterior distributions obtained from the PHN and HH datasets, as well as the union of these two posterior distributions. Two sampling strategies are considered: variable, where the time between each observation is allowed to vary, and fixed.

Under the constraint of equal observation intervals, and restricted to whole days any sampling interval between 9 days and 11 days gives a Fisher Information within 97% of the maximum for the PHN dataset. Comparatively, for the HH dataset, any sampling interval between 21 days and 28 days gives a Fisher Information within 97% of the maximum. Combining the two posterior distributions, any sampling interval between 21 and 28 days is within 97% of the maximum. However, it should be noted that a sampling interval of 23.4 days achieves only 30% of the maximum Fisher information possible in the PHN dataset, but 99% of the maximum in the HH dataset. This highlights the importance of specifying the optimal sampling strategy according to the specific scenario.

Interestingly, the optimal design spacing for the fixed strategy is *not* the minimum of the optimal design spacing for the variable strategy. We propose the following hypothesis for this phenomenon: when the observation interval is allowed to vary, we can effectively ‘spend’ a single observation close to the previous in order to potentially gain a lot of information. However, in the fixed interval strategy, this option is not available, and so to avoid ‘wasting’ observations, a more conservative strategy becomes the optimal.

To understand the difference in the optimal sampling times, recall that the expression in Equation (5) maximises the Fisher Information, which through the Cramer–Rao lower bound, can be thought of as minimising the variance of the parameter estimates [6]. This estimate inherently depends on the underlying parameters of the system: when events (i.e., infection and recovery) are happening slowly (i.e., low prevalence) then sampling should happen less often, while when events are happening frequently (i.e., high prevalence), then sampling should happen more often. In the case where little prior information about the system is available, then it may be more appropriate to adopt a ‘conservative’ sampling strategy, which here is the faster of the two presented strategies. Doing this yields a Fisher Information of 53% of the maximum for the HH dataset. The conservative strategy is presented in Appendix D. Overall, the conservative strategy generally gives good estimation accuracy (up to 10% error in a simulation-estimation study), and so is a viable ‘catch-all’ strategy in the absence of prior information such as the prevalence.

## 6 Conclusion

We have provided the first model-based estimates for the duration of a skin sore infection (between 12 and 20 days), the force of infection and basic reproductive ratio (1.1 to 1.6) in three different settings. Furthermore, the optimal sampling interval for future strategies has been determined, assuming that a study’s primary goal is to estimate the force of infection and duration of infectiousness.

Previous work on the duration of skin sore infection has calculated the lifetime of a single sore on an individual under observed treatment to be less than 7 days [4]. However, the lifetime of a single sore is unlikely to be the same as the period for which an individual is infectious, due to the presence of multiple sores on a single individual. By performing the estimation in a modelling framework, the interval-censored nature of the data has been incorporated. Although the frequentist version of this estimation technique has been utilised in other disease settings [14, 17], to our knowledge this is the first time these quantities have been computed for skin sores.

These results have been calculated using a linearised SIS model, in which the transmission rate has been assumed to be constant, and disease dynamics are at equilibrium. This assumption has allowed some simple analytic results which are often not able to be determined for traditional infectious disease dynamic models. However, it is important to note that the assumption of equilibrium dynamics is likely to be violated in real-world settings, particularly in the event of mass drug administration. Mass drug administration has been implemented in these communities in the past [1, 20], and was ongoing during the period of data collection in the RC dataset, although skin sores was not the primary outcome of the program in the RC setting [5]. It is also important to note that the SIS model structure, by construction, does not incorporate any period of immunity, or other potential disease states. Carriage (i.e. infected but not showing symptoms), in particular has been demonstrated for skin sore infections in the past [25, 11] and inclusion of carriage in models has been shown to substantially change intervention outcomes [7]. However, without longitudinal carriage data related to skin infection, or clear evidence of the contribution of carriage to infectiousness, quantifying the impact in these settings is challenging.

In the populations in which these data were collected, treatment is routinely administered for skin sores. Thus, this estimate of the infectious period is influenced by treatment, and so is likely to be lower than the *natural* infectious period. Quantification of the impact of treatment is an important research question. Within these studies, diagnosis of skin infections was actively sought and treatment recommended. However, actual uptake of treatment was not recorded. Nonetheless, treatment was probably more common than outside of the study context, meaning that our estimate of the infectious period is likely an underestimate of the infectious period outside of the study context.

There are a number of key differences between the three datasets considered. The PHN dataset only has observations of children under five years of age. Extrapolation from this dataset to the entire population should be performed with caution as the prevalence of skin sores appears to be age-specific [35], although the relative similarity of the estimates of the infectious period from the HH data (in which the general population was studied) does provide some reassurance of the estimated numbers. Further, sampling times in the PHN dataset were not fixed in advance, but were rather driven by patients or health professionals. It has been assumed these sampling times are ignorable, but further investigation into this assumption may be warranted.

As well as estimation of key parameters for models of skin sores transmission, information about future experimental designs has also been provided. Although the optimal sampling interval is a function of both the force of infection and the infectious period, being able to calculate this interval provides helpful information to improve the efficiency of future study designs, or evaluation of disease control programs.

These parameter estimates unlock future model-based investigations for skin sores. By providing estimates for both the force of infection and the duration of infectiousness, more complex models which include covariates such as scabies, non-homogenous contact patterns, and population mobility can be considered, and the impact of treatment strategies in these settings can be evaluated. It is our hope that these models will lead to the development of innovative disease control measures, the application of which will reduce the burden of skin disease and health inequalities.

## Data accessibility

The observation times for individuals, fitted posterior distributions for each dataset, and the optimisation algorithm are given in the supplementary material. Simulation algorithms and estimation methods are provided in the TMI package (https://github.com/MikeLydeamore/TMI).

## Authors’ Contributions

M.J.L., P.T.C., J.M., S.Y.C. and J.M.M. conceived the study. M.J.L. and D.J.P. designed and performed the analysis. A.J.M., W.C., Y.W., J.R.C., R.M.A. and M.I.M. provided and manipulated the data. M.J.L. wrote the manuscript. M.J.L., P.T.C., D.J.P., Y.W., A.J.M., W.C., J.R.C., J.M., S.Y.C., J.M.M. edited the manuscript. All authors gave final approval for publication.

## Acknowledgements

We would like to thank the Menzies School for Health Research and related project staff, for providing the data from the East Arnhem Healthy Skin Project, staff at the primary health care centres and the members of the remote indigenous communities for their participation. We acknowledge our partners in this work: Northern Territory Remote Health, Aboriginal Medical Services Alliance Northern Territory, Northern Territory Centre for Disease Control, One Disease and Miwatj Health and the NHMRC funded HOT NORTH initiative. We acknowledge the Lowitja Institute and the Cooperative Research Centre for Aboriginal Health who originally funded and lent significant support to the East Arnhem Healthy Skin Project.

## A MCMC Diagnostics

Here information relating to the diagnostics of the MCMC procedure is provided. Figure 7(A) shows that the posterior distributions from the PHN data have converged far from the prior distributions (which were 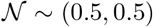), and Figure 7(B) shows that the chains are well mixed. The same conclusion can be drawn from Figure 8 for the HH dataset and for the RC dataset in Figure 9.

**Figure 7:**
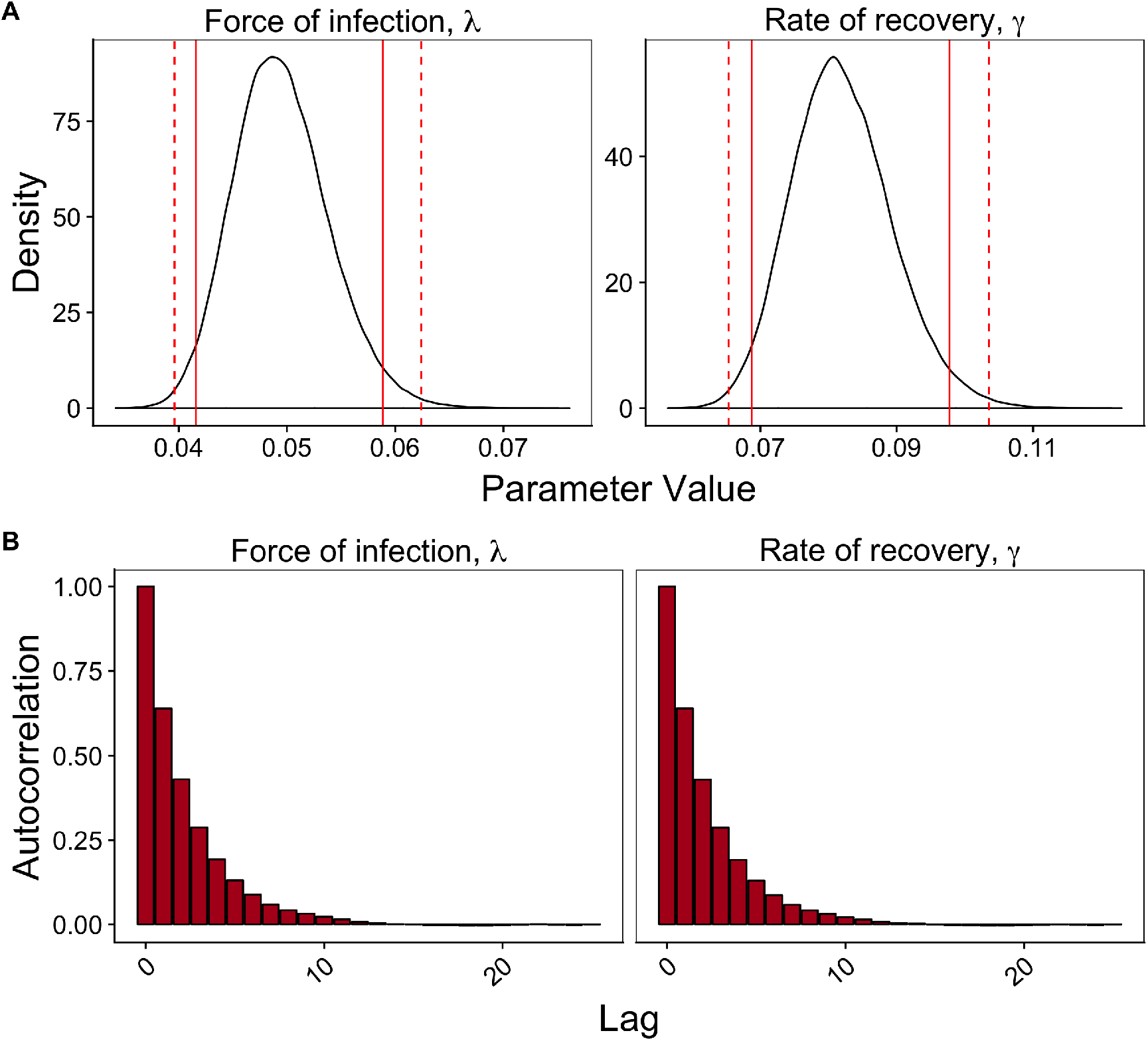
MCMC diagnostics for the PHN dataset. (A): Posterior density estimates of the force of infection, λ, and the rate of recovery, *γ*. The solid red line is a 95% credible interval, the dashed line a 99% credible interval. (B): Autocorrelation plots of the parameter values, for each of the chains.

**Figure 8:**
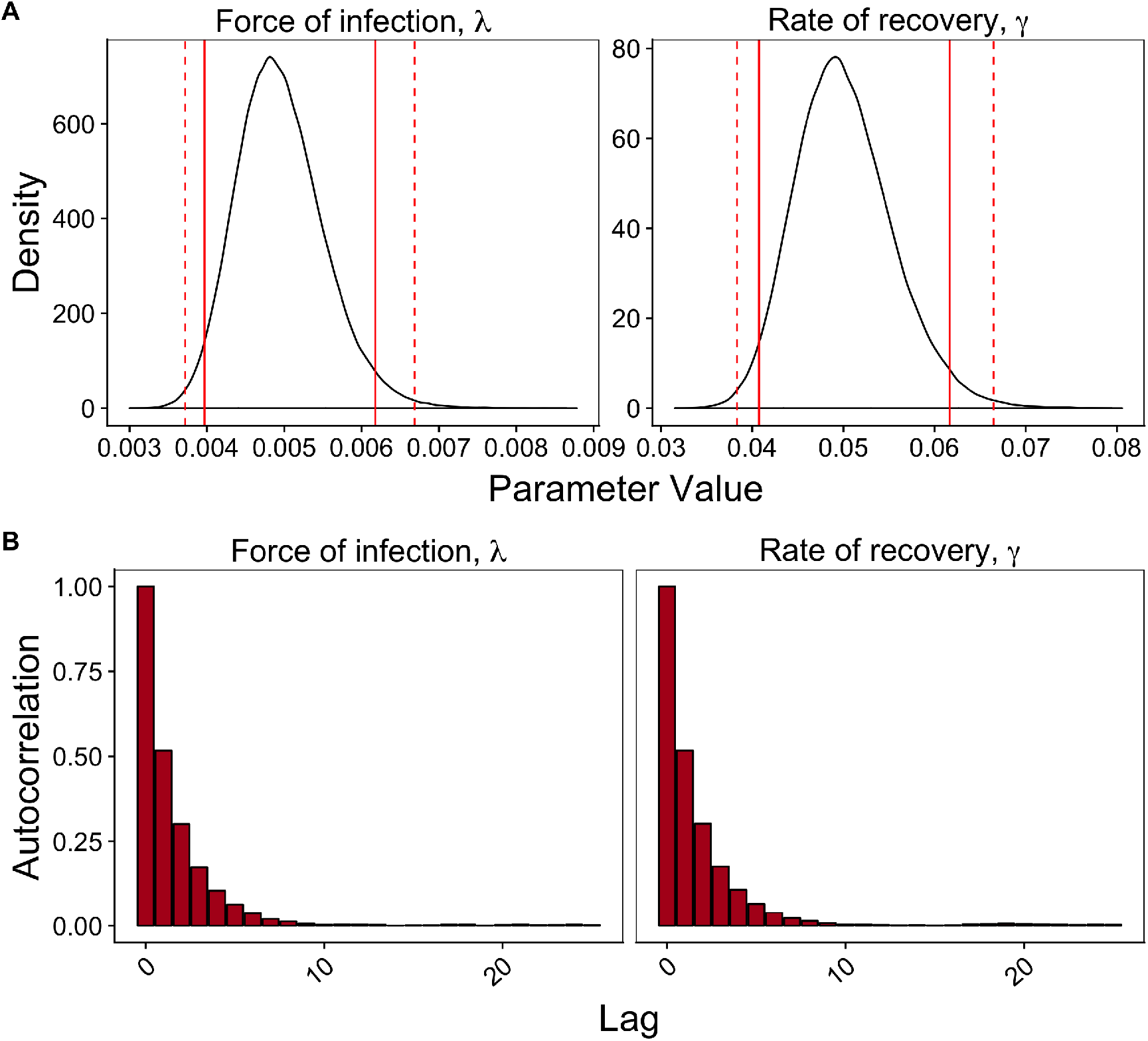
MCMC diagnostics for the HH dataset. (A): Posterior density estimates of the force of infection, λ, and the rate of recovery, *γ*. The solid red line is a 95% credible interval, the dashed line a 99% credible interval. (B): Autocorrelation plots of the parameter values, for each of the chains.

**Figure 9:**
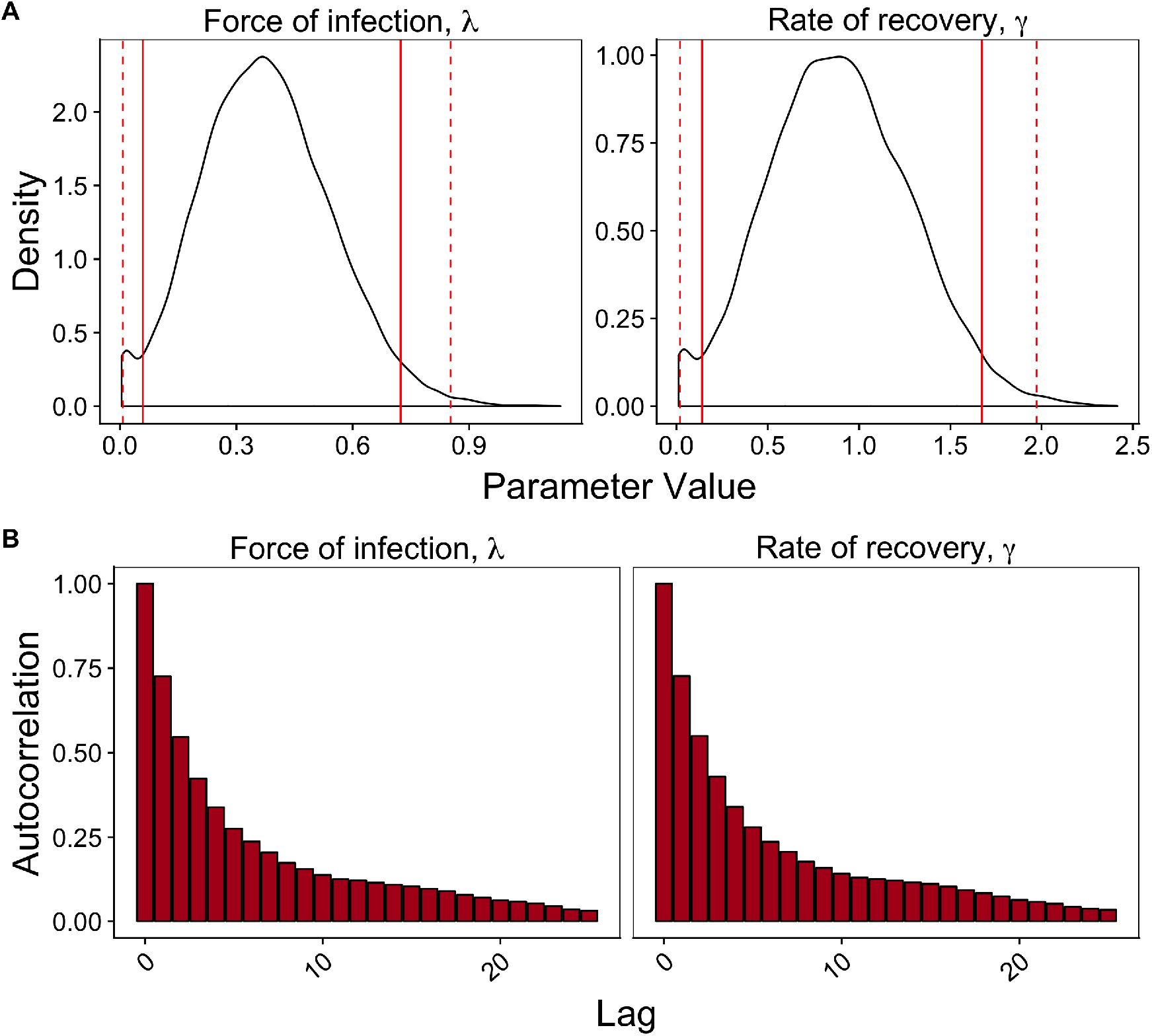
MCMC diagnostics for the RC dataset. (A): Posterior density estimates of the force of infection, λ, and the rate of recovery, *γ*. The solid red line is a 95% credible interval, the dashed line a 99% credible interval. (B): Autocorrelation plots of the parameter values, for each of the chains.

## B Derivation of the Fisher Information matrix

The Fisher Information matrix is a representation of the amount of information that is contained in a model with parameters ***θ***, about some observable value. The Fisher Information matrix is defined as,

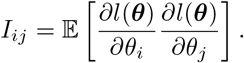

Under some regularity conditions (which are assumed to be true), this is equivalent to,

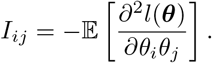

For the linearised SIS model, the Fisher Information matrix can be analytically determined and evaluated rapidly for a wide range of values for the time between each observation, ***δ***. Here, only the case of a single individual is considered, but note that extension to *N* individuals simply results in the Fisher Information being multiplied by *N*, as there is an assumption that all individuals are identical.

Define the function,

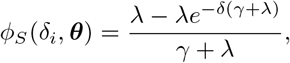

to be the probability that an individual is infected at time *δ_i_*, given they were susceptible at time 0. Similarly, define,

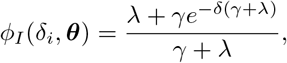

to be the probability that an individual is infected at time *δ_i_*, given they were infected at time 0. For convenience, supress the dependence on ***θ*** while deriving the Fisher Information matrix.

The likelihood function in Equation (4) can then be expressed as

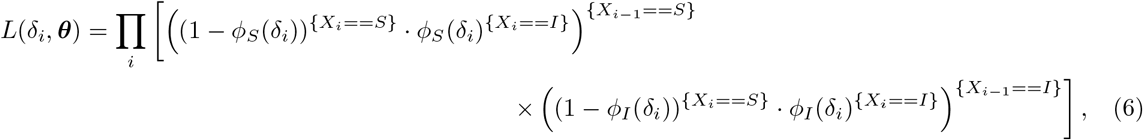

where {*X_i_* == *S*} represents an indicator function. Taking the log of Equation (6) gives

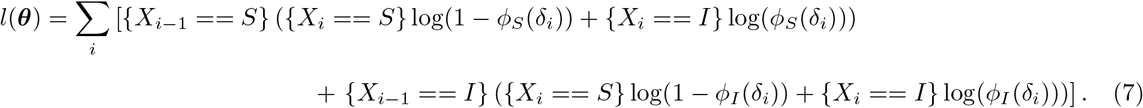

The only terms of Equation (7) that contain ***θ*** are the functions *ϕ_S_*(*δ_i_*) and *ϕ_I_*(*δ_i_*), and the log likelihood is linear in these functions. As such, the second partial derivatives of Equation (7) are simply

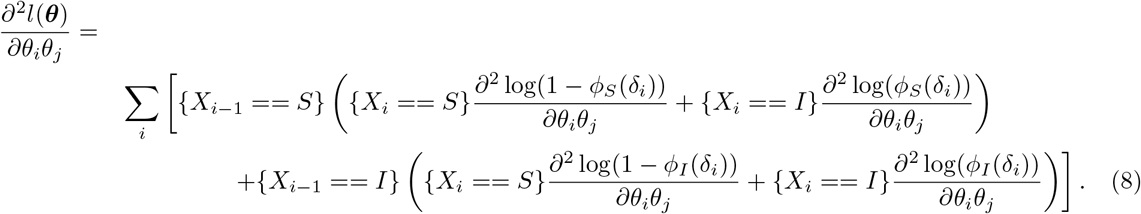

The next step to determine the Fisher Information matrix is to consider the expectation of the product of the two random variables. In this case, *X_i_* is Bernoulli, with probability of success (infection) of either *φ_S_*(*δ_j_*) if *X*_*i*−1_ = *S* or *ϕ_I_*(*δ_i_*) if *X*_*i*−1_ = *I*. Using the law of total probability, it follows that

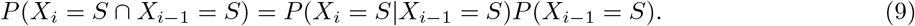

The first term of Equation (9) is simply the probability of failure (that is, not infected), given an individual was previously susceptible, which is (1 – *ϕ_S_*). To calculate the second term, recall that the time between the *i*th and the (*i* + 1)th observation is *δ_i_*. Consider a discrete time Markov chain, with probability matrix

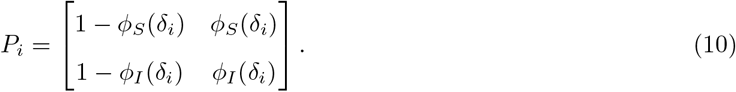

Then,

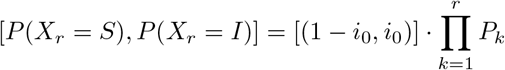

represents the probability that an individual is susceptible or infected at observation *r*, where *i*_0_ represents the probability that an individual is initially infected, given here by the prevalence, *I*^*^. For convenience, define *p_s,r_* to be the first element of this vector, and *p_i,r_* = 1 – *p_s,r_* to be the second element.

As each *X_i_* is Bernoulli, it follows from Equation (9) that the joint expectation is

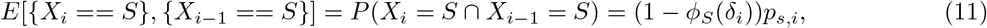

and the other forms of this expectation follow similarly.

Finally, taking the expectation of Equation (8) gives

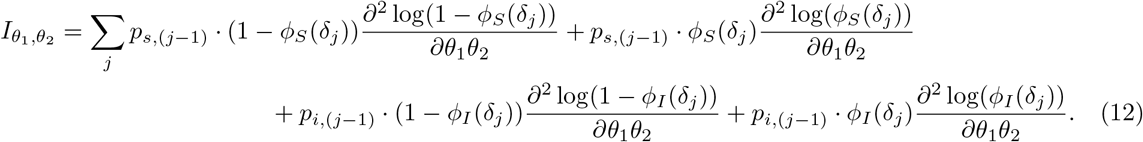

The expressions for each second derivative were found using Sage, and are implemented for numeric evaluation as part of the `TMI` package (https://github.com/MikeLydeamore/TMI).

### B.1 Constant time between observations

If it is assumed that the time between observations is constant, that is,

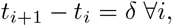

then the *P* matrix in Equation (10) is independent of *i*. Because of this, an analytic expression for *P^r^* is [38],

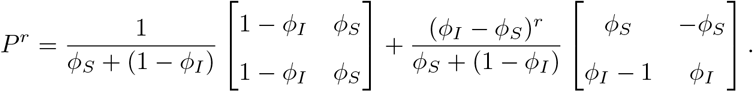

The expression for the Fisher Information in Equation (12) is verified in the constant time between observations case again through a simulation estimation approach. For a given set of parameters, ***θ*** = {λ, *γ*}, 64 populations of the SIS model are simulated, and parameters estimated under each sample spacing, *δ* ∈ {5,10,…, 40}. The determinant of the covariance matrix under each sample spacing is calculated and normalised. The results of this verification are shown in Figure 10.

**Figure 10:**
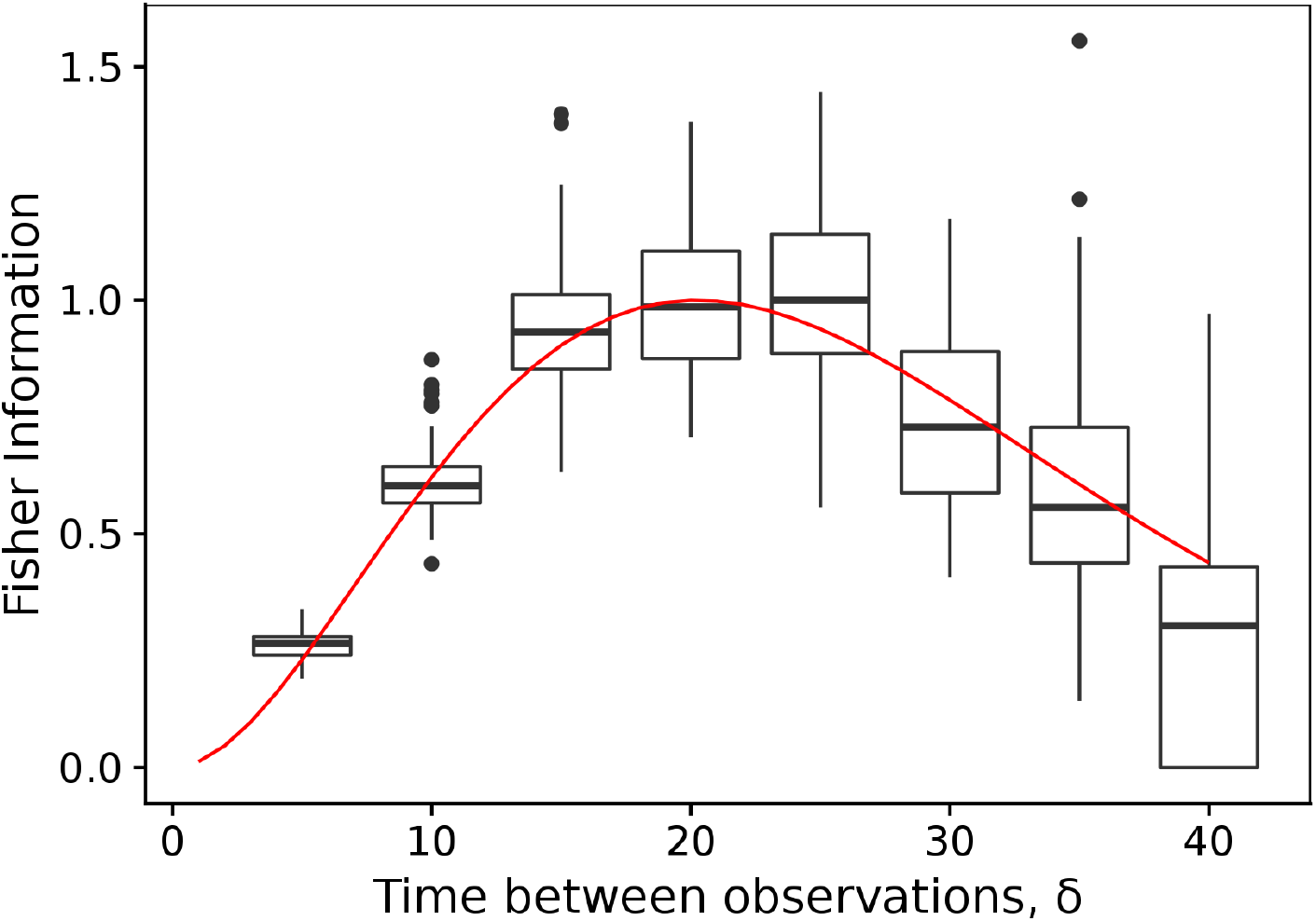
Numeric verification of the expressions for the Fisher Information matrix. The boxplot represents the summary of the determinants of the covariance matrix, and the red line the analytic expression for the Fisher Information matrix, with each curve normalised separately. The chosen parameters were λ = 1/60 and *γ* = 1/20.

## C Optimal sampling strategy diagnostics

We utilise the induced natural selection heuristic for finding optimal sampling strategies [32]. We choose the following parameters:

- Initial designs, *D* ~ *U*[1,40], with |*D*| = 2000,
- Number of generations, *W* = 50,
- Pertubation function, 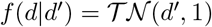,
- Acceptance criteria: retain top 40, 30, 20, 10 and 5 designs (10 times each),
- Newly sampled designs, *m*: 5, 10, 20, 30 and 40 designs (10 times each).

Presented here are diagnostics which give evidence of the convergence of the induced natural selection heuristic. Figures 11, 12 and 13 show that the Fisher information matrix has converged (panel A), and that the chosen values for the sampling intervals also appear to have converged (panel B) for the variable interval strategies. The same conclusion is drawn for the fixed interval strategies, as shown in Figurse 14, 15 and 16.

**Figure 11:**
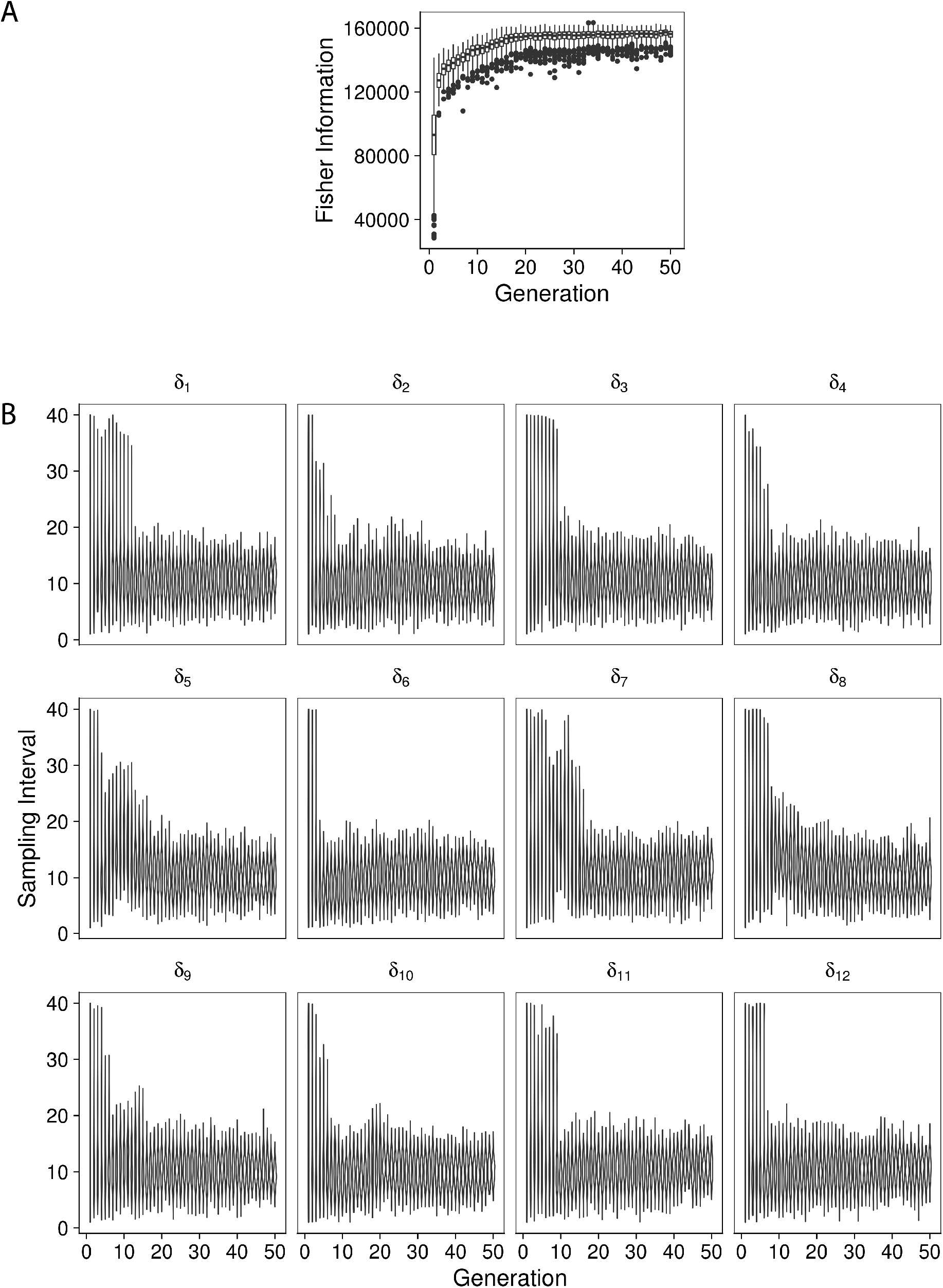
Accepted values of the sampling intervals for the PHN dataset with a variable sampling interval, as a function of the generation of the estimation algorithm. The space appears well explored, and the solution appears converged after 50 generations.

**Figure 12:**
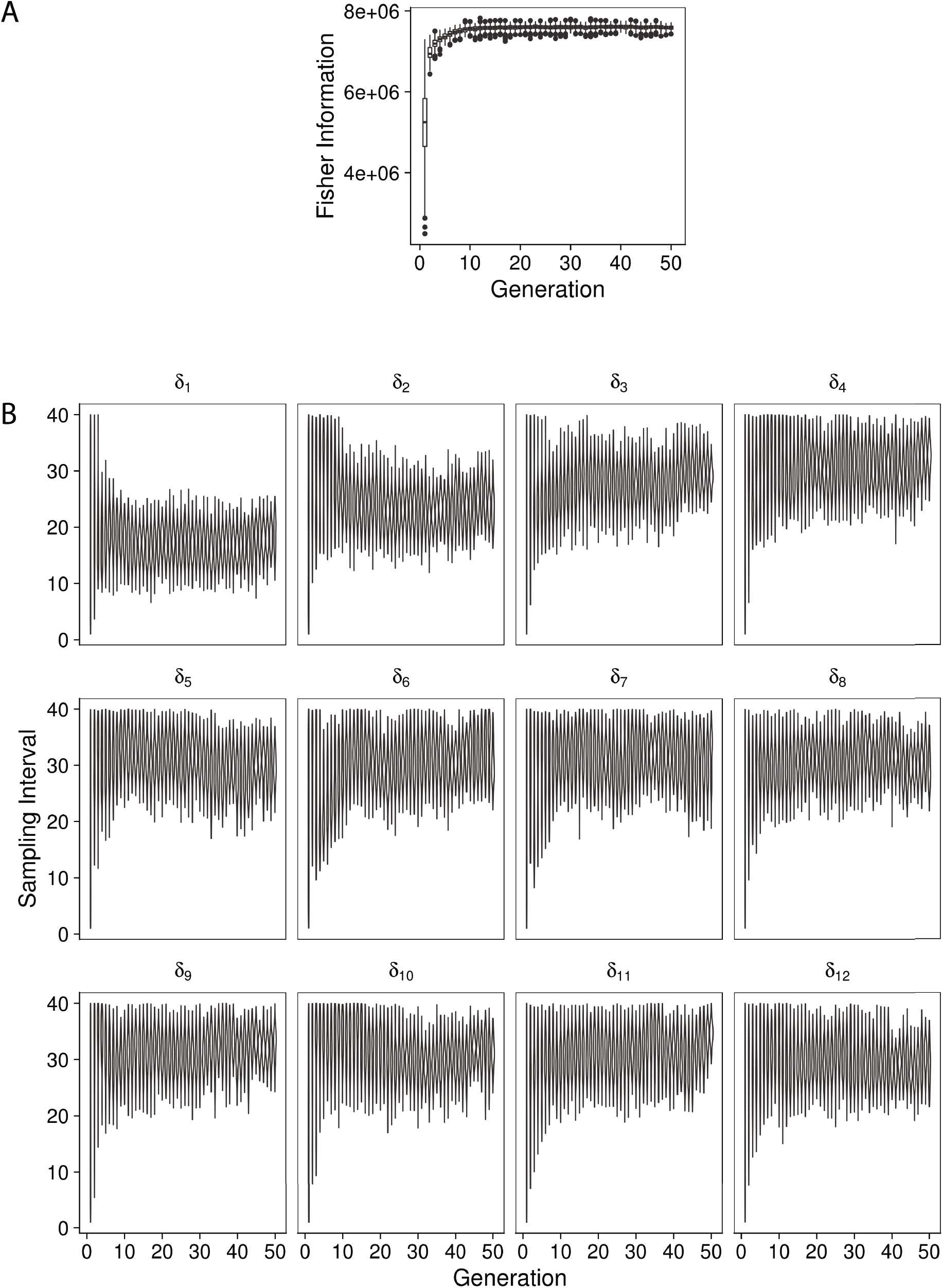
Accepted values of the sampling intervals for the HH dataset with a variable sampling interval, as a function of the generation of the estimation algorithm. The space appears well explored, and the solution appears converged after 50 generations.

**Figure 13:**
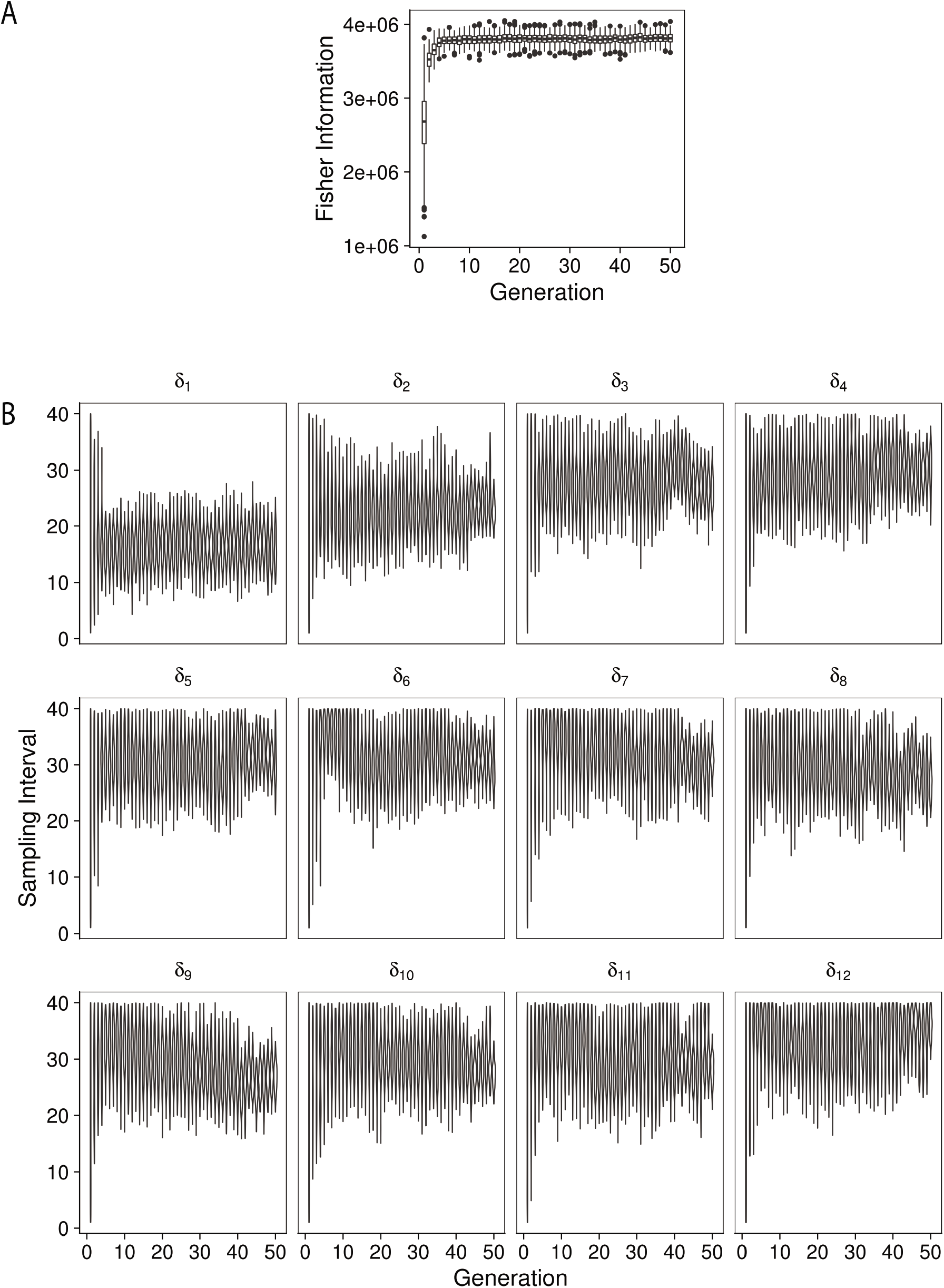
Accepted values of the sampling intervals for the combined dataset with a variable sampling interval, as a function of the generation of the estimation algorithm. The space appears well explored, and the solution appears converged after 50 generations.

**Figure 14:**
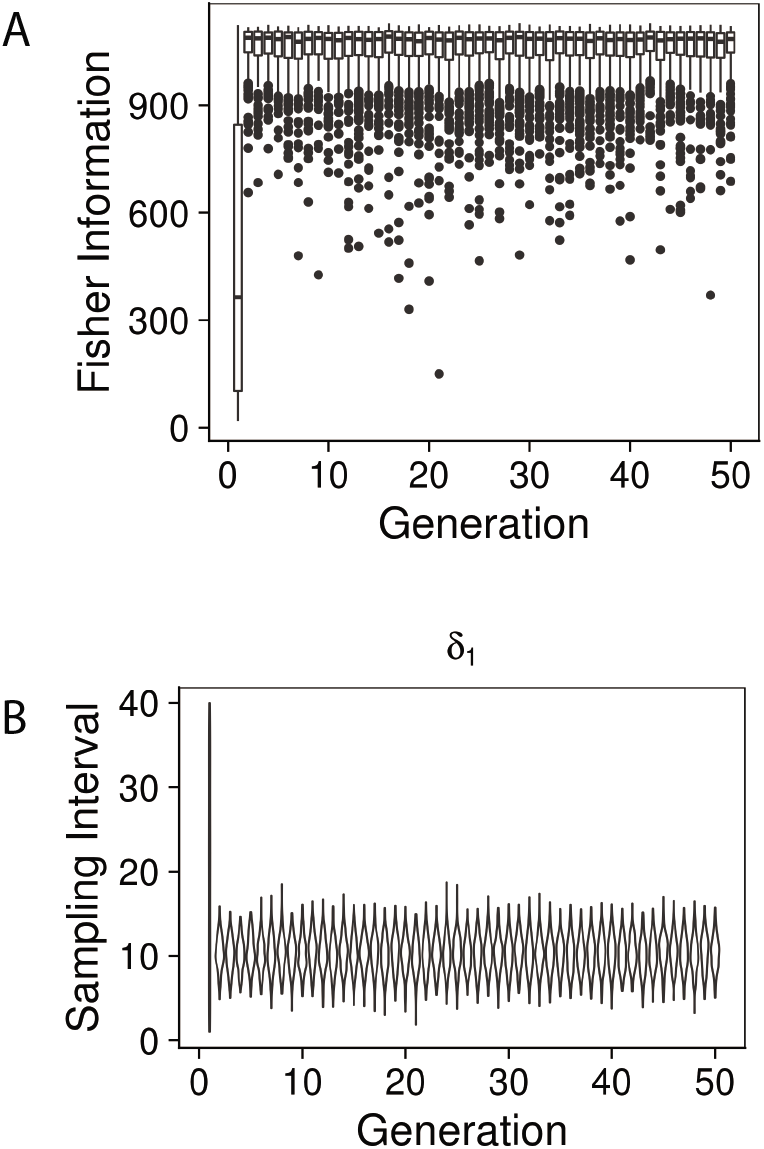
Accepted values of the sampling intervals for the PHN dataset with a fixed sampling interval, as a function of the generation of the estimation algorithm. The space appears well explored, and the solution appears converged after 50 generations.

**Figure 15:**
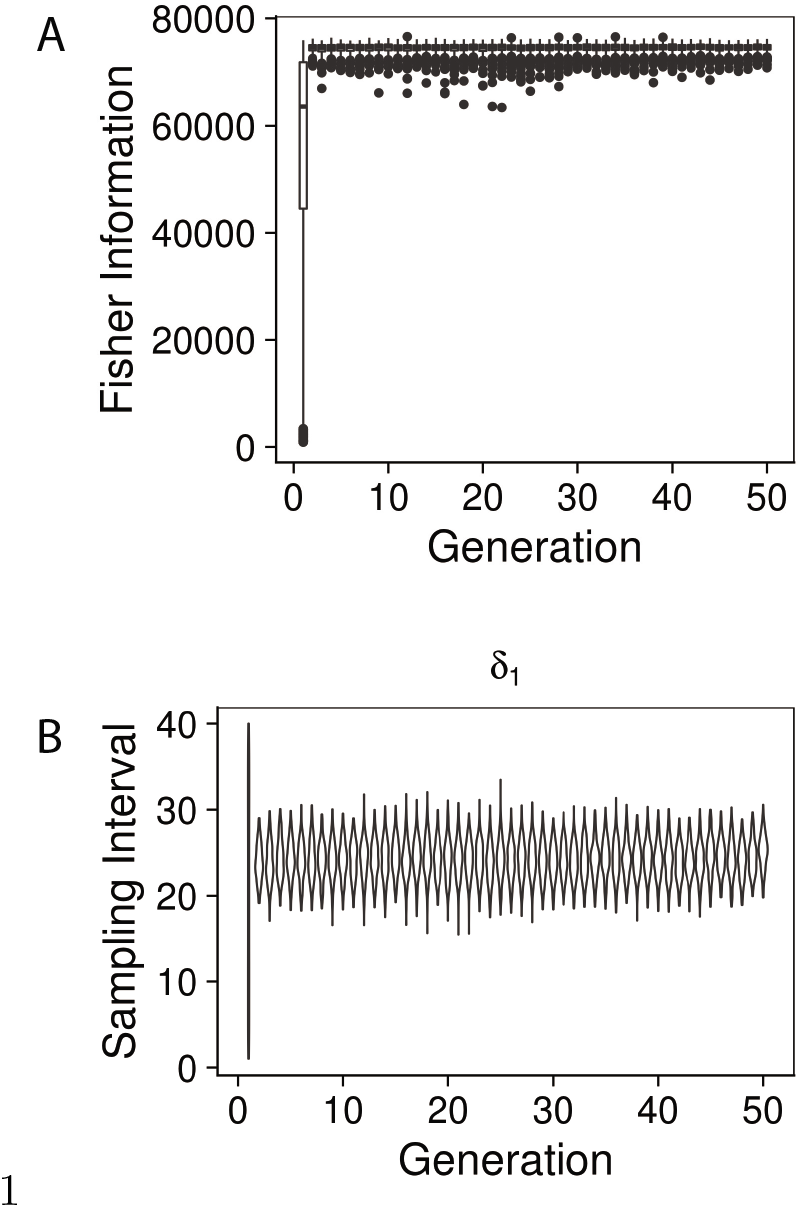
Accepted values of the sampling intervals for the HH dataset with a fixed sampling interval, as a function of the generation of the estimation algorithm. The space appears well explored, and the solution appears converged after 50 generations.

**Figure 16:**
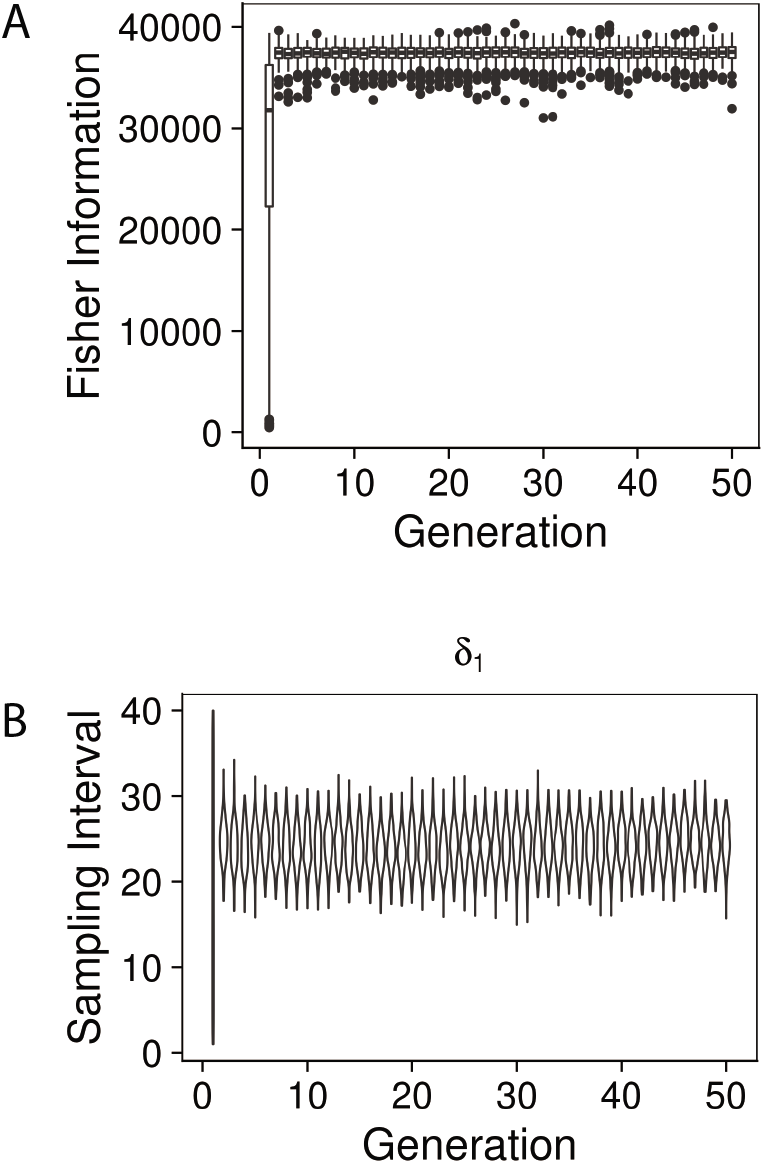
Accepted values of the sampling intervals for the combined dataset with a fixed sampling interval, as a function of the generation of the estimation algorithm. The space appears well explored, and the solution appears converged after 50 generations.

## D Conservative sampling strategy

In Section 5, it was found that the optimal sampling strategy favoured a longer sampling interval in the absence of prevalence information. While this performed well for the relatively low prevalence observed in the setting where the HH dataset was collected, the strategy performed poorly in the higher prevalence setting where the PHN dataset was collected. A potential alternative strategy would be the *conservative* strategy, where the population is sampled every 10.45 days (as per the high prevalence setting). This strategy favours oversampling at the penalty of potentially broader credible intervals for parameter estimates in a low prevalence setting. A simulation estimation study of this, with 20 observations, is shown in Figure 17. The conservative strategy appears to perform well in the low-prevalence setting, although the variability in estimates is high (up to a 10% estimate error, compared with a 6% estimate error in the high prevalence setting). As this sampling strategy is optimal for the high prevalence setting, this study suggests that the conservative strategy may be a valid ‘catch-all’ strategy.

**Figure 17:**
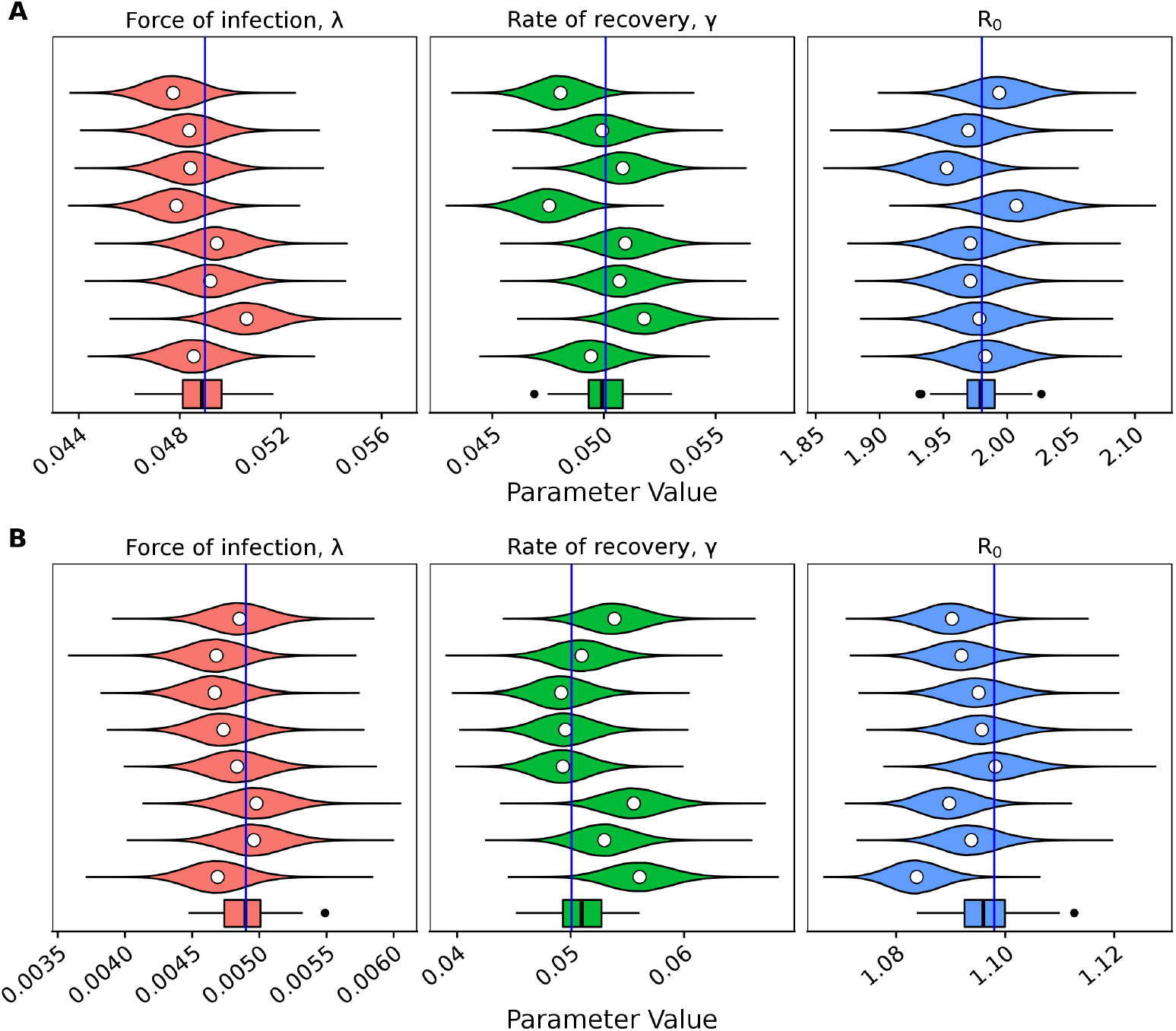
Marginal posterior distributions for the force of infection, λ, the rate of recovery *γ*, and the basic reproductive ratio, *R*_0_, from 8 randomly generated populations from the linearised SIS model under the conservative sampling strategy (every 10.45 days, for 20 successive observations) under (A) a high prevalence setting, and (B) a low prevalence setting. The mean of each distribution is given by the white circle. The boxplot at the bottom of each panel represents the means of 64 marginal posteriors. The true value which was used to generate each population is represented by the blue line (high prevalence λ = 0.049, low prevalence λ = 0.0049, *γ* = 1/19.97)

## Bibliography

1 R. M. Andrews, T. Kearns, C. Connors, C. Parker, K. Carville, B. J. Currie, and J. R. Carapetis. A regional initiative to reduce skin infections amongst Aboriginal children living in remote communities of the Northern Territory, Australia. PLoS Neglected Tropical Diseases, 3(11):e554, 2009.

2 A. J. Black, N. Geard, J. M. McCaw, J. McVernon, and J. V. Ross. Characterising pandemic severity and transmissibility from data collected during first few hundred studies. Epidemics, 19(Supplement C):61–73, June 2017.

3 A. C. Bowen, A. Mahé, R. J. Hay, R. M. Andrews, A. C. Steer, S. Y. C. Tong, and J. R. Carapetis. The global epidemiology of impetigo: A systematic review of the population prevalence of impetigo and pyoderma. PLOS ONE, 10(8):e0136789, 2015.

4 A. C. Bowen, S. Y. C. Tong, R. M. Andrews, I. M. O’Meara, M. I. McDonald, M. D. Chatfield, B. J. Currie, and J. R. Carapetis. Short-course oral co-trimoxazole versus intramuscular benzathine benzylpeni-cillin for impetigo in a highly endemic region: An open-label, randomised, controlled, non-inferiority trial. The Lancet, 384(9960):2132–2140, 2014.

5 J. R. Carapetis, C. Connors, D. Yarmirr, V. Krause, and B. J. Currie. Success of a scabies control program in an Australian Aboriginal community. The Pediatric Infectious Disease Journal, 6:494–499, 1997.

6 G. Casella and R. L. Berger. Statistical Inference, volume 2. Duxbury Pacific Grove, CA, 2002.

7 R. H. Chisholm, P. T. Campbell, Y. Wu, S. Y. C. Tong, J. McVernon, and N. Geard. Implications of asymptomatic carriers for infectious disease transmission and control. Open Science, 5(2):172341, 2018.

8 D. B. Clucas, K. S. Carville, C. Connors, B. J. Currie, J. R. Carapetis, and R. Andrews. Disease burden and health-care clinic attendances for young children in remote Aboriginal communities of northern Australia. Bulletin of the World Health Organization, 86(4):275–281, 2008.

9 J. N. Cole, T. C. Barnett, V. Nizet, and M. J. Walker. Molecular insight into invasive group A streptococcal disease. Nature Reviews Microbiology, 9(10):724–736, Oct. 2011.

10 A. S. Dajani, P. Ferrieri, and L. W. Wannamaker. Natural history of impetigo: II. Etiologic agents and bacterial interactions. Journal of Clinical Investigation, 51(11):2863–2871, 1972.

11 B. A. Dudding, J. W. Burnett, S. S. Chapman, and L. W. Wannamaker. The role of normal skin in the spread of Streptococcal pyoderma. The Journal of Hygiene, 68(1):19–28, 1970.

12 P. Ferrieri, A. S. Dajani, L. W. Wannamaker, and S. S. Chapman. Natural history of impetigo. I. Site sequence of acquisition and familial patterns of spread of cutaneous streptococci. The Journal of Clinical Investigation, 51(11):2851–2862, 1972.

13 R. A. Fisher. The Design of Experiments. Oliver & Boyd, 1935.

14 N. C. Grassly, M. E. Ward, S. Ferris, D. C. Mabey, and R. L. Bailey. The natural history of trachoma infection and disease in a Gambian cohort with frequent follow-up. PLOS Neglected Tropical Diseases, 2(12):e341, 2008.

15 J. Gruger, R. Kay, and M. Schumacher. The validity of inferences based on incomplete observations in disease state models. Biometrics, 47(2):595–605, 1991.

16 N. D. Hysmith, E. L. Kaplan, P. P. Cleary, D. R. Johnson, T. A. Penfound, and J. B. Dale. Prospective longitudinal analysis of immune responses in pediatric subjects after pharyngeal acquisition of Group A Streptococci. Journal of the Pediatric Infectious Diseases Society, 6(2):187–196, 2017.

17 R. Ivanek, J. Osterberg, R. Gautam, and S. S. Lewerin. Salmonella fecal shedding and immune responses are dose- and serotype-dependent in pigs. PLOS ONE, 7(4):e34660, 2012.

18 C. Jackson. Multi-State Models for Panel Data: The msm Package for R. Journal of Statistical Software, 38(8):28, 2011.

19 T. Kearns, D. Clucas, C. Connors, B. J. Currie, J. R. Carapetis, and R. M. Andrews. Clinic Attendances during the first 12 months of life for Aboriginal children in five remote communities of northern Australia. PLOS ONE, 8(3):1–5, 2013.

20 T. M. Kearns, R. Speare, A. C. Cheng, J. McCarthy, J. R. Carapetis, D. C. Holt, B. J. Currie, W. Page, J. Shield, R. Gundjirryirr, L. Bundhala, E. Mulholland, M. Chatfield, and R. M. Andrews. Impact of an Ivermectin mass drug administration on scabies prevalence in a remote Australian Aboriginal community. PLOS Neglected Tropical Diseases, 9(10):e0004151, 2015.

21 M. J. Keeling and P. Rohani. Modeling Infectious Diseases in Humans and Animals. Princeton University Press, 2008.

22 W. O. Kermack and A. G. McKendrick. A contribution to the mathematical theory of epidemics. Proceedings of the Royal Society of London A: Mathematical, Physical and Engineering Sciences, 115:700–721, 1927.

23 I. M. Longini, J. S. Koopman, M. Haber, and G. A. Cotsonis. Statistical inference for infectious diseases: Risk-specific household and community transmission parameters. American Journal of Epidemiology, 128(4):845–859, Oct. 1988.

24 J. S. Maddox, J. C. Ware, and H. C. Dillon. The natural history of streptococcal skin infection: Prevention with topical antibiotics. Journal of the American Academy of Dermatology, 13(2):207–212, 1985.

25 M. I. McDonald, R. J. Towers, R. M. Andrews, N. Benger, B. J. Currie, and J. R. Carapetis. Low rates of Streptococcal pharyngitis and high rates of pyoderma in Australian Aboriginal communities where acute rheumatic fever is hyperendemic. Clinical Infectious Diseases, 43(6):683–689, 2006.

26 E. McMeniman, L. Holden, T. Kearns, D. B. Clucas, J. R. Carapetis, B. J. Currie, C. Connors, and R. M. Andrews. Skin disease in the first two years of life in Aboriginal children in East Arnhem Land. Australasian Journal of Dermatology, 52(4):270–273, 2011.

27 R. Moss, A. Zarebski, P. Dawson, and J. M. McCaw. Forecasting influenza outbreak dynamics in Melbourne from Internet search query surveillance data. Influenza and other respiratory viruses, 10(4):314–323, 2016.

28 E. O. Nsoesie, J. S. Brownstein, N. Ramakrishnan, and M. V. Marathe. A systematic review of studies on forecasting the dynamics of influenza outbreaks. Influenza and Other Respiratory Viruses, 8(3):309–316, May 2014.

29 P. O’Neill and G. Roberts. Bayesian inference for partially observed stochastic epidemics. Journal of the Royal Statistical Society: Series A (Statistics in Society), 162(1):121–129, 1999.

30 C.-l. Y. Ong, C. M. Gillen, T. C. Barnett, M. J. Walker, and A. G. McEwan. An antimicrobial role for Zinc in innate immune defense against Group A Streptococcus. The Journal of Infectious Diseases, 209(10):1500–1508, May 2014.

31 D. Pagendam and P. Pollett. Locally optimal designs for the simple death process. Journal of Statistical Planning and Inference, 140(11):3096–3105, 2010.

32 D. J. Price, N. G. Bean, J. V. Ross, and J. Tuke. An induced natural selection heuristic for finding optimal Bayesian experimental designs. Computational Statistics & Data Analysis, 126:112–124, Oct. 2018.

33 L. Pronzato and E. Walter. Robust experiment design via stochastic approximation. Mathematical Biosciences, 75(1):103–120, 1985.

34 M. J. Rodríguez-Ortega, N. Norais, G. Bensi, S. Liberatori, S. Capo, M. Mora, M. Scarselli, F. Doro, G. Ferrari, I. Garaguso, T. Maggi, A. Neumann, A. Covre, J. L. Telford, and G. Grandi. Characterization and identification of vaccine candidate proteins through analysis of the group A *Streptococcus* surface proteome. Nature Biotechnology, 24(2):191–197, Feb. 2006.

35 L. Romani, J. Koroivueta, A. C. Steer, M. Kama, J. M. Kaldor, H. Wand, M. Hamid, and M. J. Whitfeld. Scabies and impetigo prevalence and risk factors in Fiji: A national survey. PLOS Neglected Tropical Diseases, 9:e0003452, 2015.

36 L. Romani, A. C. Steer, M. J. Whitfeld, and J. M. Kaldor. Prevalence of scabies and impetigo worldwide: A systematic review. The Lancet Infectious Diseases, 15(8):960–967, 2015.

37 Stan Development Team. RStan: The R interface to Stan. 2018. R package version 2.17.3.

38 E. A. Suess and B. E. Trumbo. Introduction to Probability Simulation and Gibbs Sampling with R. Use R! Springer, New York, 2010. OCLC: ocn620103481.

39 M. J. Walker, J. D. McArthur, F. McKay, and M. Ranson. Is plasminogen deployed as a Streptococcus pyogenes virulence factor? Trends in Microbiology, 13(7):308–313, July 2005.

40 A. E. Zarebski, P. Dawson, J. M. McCaw, and R. Moss. Model selection for seasonal influenza forecasting. Infectious Disease Modelling, 2(1):56–70, Feb. 2017.

